# PRRC2A, PRRC2B and PRRC2C are Stress Granule Proteins that Promote Translation Through Association with the eIF3 complex

**DOI:** 10.64898/2026.02.24.707808

**Authors:** Jie Qi Huang, Eileigh Kadijk, Karl J. Schreiber, Zi Hao Liu, Rachel W. Chiang, Zhixin Zhang, Kevin Guttman, Tian Hao Huang, Sylvia M.T. Almeida, Olena Zhulyn, Alan Moses, Julie D. Forman-Kay, John L. Rubinstein, Ji-Young Youn

**Author notes:** Data deposition: Mass spectrometry data have been deposited in the Mass Spectrometry Interactive Virtual Environment (MassIVE, http://massive.ucsd.edu). MassIVE ID: MSV000100946(ftp://massive-ftp.ucsd.edu/v12/MSV000100946/) MassIVE ID: MSV000100945 (ftp://massive-ftp.ucsd.edu/v12/MSV000100945/) MassIVE ID: MSV000100944 (ftp://massive-ftp.ucsd.edu/v12/MSV000100944/).

## Abstract

Regulation of mRNA translation is essential for cellular homeostasis, and its dysregulation contributes to cancer, neurodegeneration, and developmental disorders. Stress granules are cytosolic condensates that form during stress-induced translation arrest and are enriched in mRNAs, translation factors, and RNA-binding proteins, but how stress granule proteins modulate translation remains poorly understood. Here, we identify the stress granule components Proline-Rich Coiled-Coil A, B, and C (PRRC2 proteins) as translation regulators. PRRC2 proteins are large, intrinsically disordered paralogs conserved across jawed vertebrates. Functional proteomics revealed that all PRRC2 proteins associate with the 48S translation initiation complex (PIC), whereas PRRC2B additionally interacts with nuclear proteins. Under stress, the proximal interaction network of PRRC2 proteins undergoes dynamic remodeling, including increased interactions with the stress granule scaffold G3BP1. Genetic perturbation shows that the PRRC2 proteins influence stress granule assembly in a context-specific manner, and are collectively required for cell growth in basal conditions due to their essential role in translation. Cells with reduced PRRC2 proteins exhibit a significant reduction in the abundance of more than half of the proteome, with a bias toward translational targets of eIF3d and eIF4G2. Interaction domain mapping and AlphaFold3 modeling revealed that an α helix within the putative coiled-coil domain of PRRC2C mediates interactions with the eIF3 core complex. This modeling places the PRRC2C α helix in a previously unassigned region of a published cryo-EM density map, validating the protein interaction and the mechanistic role of PRRC2C in translation control. Together, these findings establish PRRC2 proteins as components of the translation initiation machinery that regulate translation through their interactions with the eIF3 complex and other components of the 48S PIC factors, providing a direct mechanistic link between stress granule proteins and translational control.

---

Regulation of mRNA translation is fundamental to cellular function, governing processes ranging from cell growth to stress response. Under optimal conditions, efficient translation sustains biosynthetic capacity and cellular fitness. In contrast, environmental stress or cytotoxic insults trigger repression of global protein synthesis via activation of integrated stress response (ISR), which induces selective translation of mRNAs required for adaptation and survival. The dynamic reprogramming of translation is essential for maintaining cellular homeostasis, and its disruption contributes to diverse pathological states, including cancer, neurodegeneration, and developmental disorders^1–4^.

Stress granules (SGs) are biomolecular condensates intimately linked to translation control^5,6^. These cytoplasmic ribonucleoprotein assemblies form rapidly upon inhibition of translation initiation, typically downstream of ISR activation. ISR converges on the phosphorylation of eIF2⍺, which stalls translation initiation and promotes the condensation of mRNAs, RNA-binding proteins and translation factors to form SGs^7–11^. This condensation is reliant on multivalent interactions between SG components, which include transient and stable interactions^12–15^. Although SGs have historically been viewed as storage or triage sites for translationally repressed mRNAs, emerging evidence suggests more nuanced roles, including the augmentation of the ISR-activated translation programme ^16^. Therefore, there is a need to better understand the role of SG-associated proteins in translational regulation. Understanding how these proteins function under basal conditions can inform how SG proteins may regulate translation under stress.

Several lines of evidence suggest that SG proteins function beyond stress-induced condensates. Proteomic studies demonstrated extensive interaction networks among hundreds of SG proteins even in unstressed cells, despite the absence of microscopically observable granules^17,18^. In addition, multiple SG-localized proteins associate with actively translating ribosomes under basal conditions, including G3BP1, UBAP2L, FMR1 and STAU1, implicating them directly in translation regulation^19–26^.

The Proline-Rich Coiled-coil proteins PRRC2A, PRRC2B and PRRC2C (hereafter PRRC2 proteins) constitute a poorly characterized family of paralogous proteins that are strongly connected in SG-proximal interaction networks under basal conditions^17^. Genetic studies indicate that PRRC proteins are essential for mammalian development, as individual gene disruption causes preweaning lethality in mice^27,28^. Increasing evidence links PRRC2 proteins to translation regulation. All three human paralogs promote translation of mRNAs containing upstream open reading frames (uORF) or overlapping ORFs, and the *Drosophila melanogaster* ortholog *nocte* similarly enhances translation of uORF-containing transcripts, supporting evolutionary conservation of this function^29,30^. PRRC2B has additionally been implicated in translation of cell-cycle-associated mRNAs and stabilization of 5’TOP mRNAs during nutrient stress^31,32^. Consistent with these functions, PRRC2 proteins exhibit RNA-binding capacity: PRRC2A is predicted to bind to RNA^33^. The mouse paralogs Prrc2a and Prrc2b have been proposed to act as m6A readers, a modification increasingly recognized as a regulator of mRNA translation and cell fate decisions^27,34–38,38–40^.

PRRC2 proteins have been reported to assemble cotranslationally with key translation initiation factors, including eIF3b, eIF4G1 and eIF4E^29^. Canonical translation initiation begins with the formation of the 43s preinitiation complex (PIC), composed of the 40S ribosomal subunit, initiator Met-tRNA, and initiation factors, eIF1, eIF1A, eIF2, eIF3, and eIF5. Recruitment of the eIF4F cap-binding complex (eIF4E, eIF4A, eIF4G) facilitates mRNA engagement and formation of the scanning 48S pre-initiation complex (PIC), which identifies the start codon before joining with the 60S ribosomal subunit to generate an elongation-competent 80S ribosome ^3,41^.

Among the translation initiation factors, the eIF3 complex is the largest and most functionally versatile. It participates in nearly all stages of translation, including PIC recruitment, AUG scanning, start-codon recognition, reinitiation, specialized translation initiation, and ribosome recycling^42–44^. The mammalian complex comprise 13 subunits (eIF3a-m), including the eight core subunits (a,c,e,f,h,k,l, and m) that reside near the mRNA exit channel and the five peripheral subunits (b, d, g, i, and j) placed on the opposite side of the ribosome, interacting with the core via the disordered C-terminal tail of eIF3a. Importantly, several eIF3 subunits are required for efficient translation of mRNAs containing uORFs through mechanisms such as leaky scanning^45–47^, suggesting a functional intersection between PRRC2 proteins and eIF3-mediated translational control.

Here, we combine cell biology, genetics and functional proteomics to define the redundant and divergent functions of the PRRC2 protein family. We show that PRRC2 proteins share function as translation regulators required for optimal cell growth in human cells. We further uncover a divergent nuclear role for PRRC2B, suggesting a connection between SG-associated factors and nuclear biological processes. Finally, using PRRC2C as a representative paralog, we map a direct interaction with the eIF3 complex, mediated by an α-helical region within its predicted coiled-coil region, establishing PRRC2C as a previously unrecognized translation initiation factor.

## RESULTS

### PRRC2A, PRRC2B, and PRRC2C proteins, central in stress granule proximal interaction network, are intrinsically disordered proteins with shared and distinct molecular features

Previous work reported a high-resolution proximity map of RNA granules, including SGs and a related RNA-rich compartment called P-bodies, using a proximity-dependent biotinylation technique called BioID^17^. Leveraging an abortive biotin ligase enzyme (e.g., BirA*, or its molecularly evolved versions, miniTurbo) fused to a protein of interest (*i.e.,* bait) that biotinylates nearby proteins within ∼20 nm range in living cells (**Fig. 1A**), BioID reports proximal interactions, providing the local molecular neighbourhood of the bait in its native cellular context ^48–50^. BioID profiling of ∼120 proteins associated with mRNA life cycle, SGs and P-bodies revealed a dense network of pre-existing protein interactions among SG proteins under no-stress conditions.

**Figure 1.**
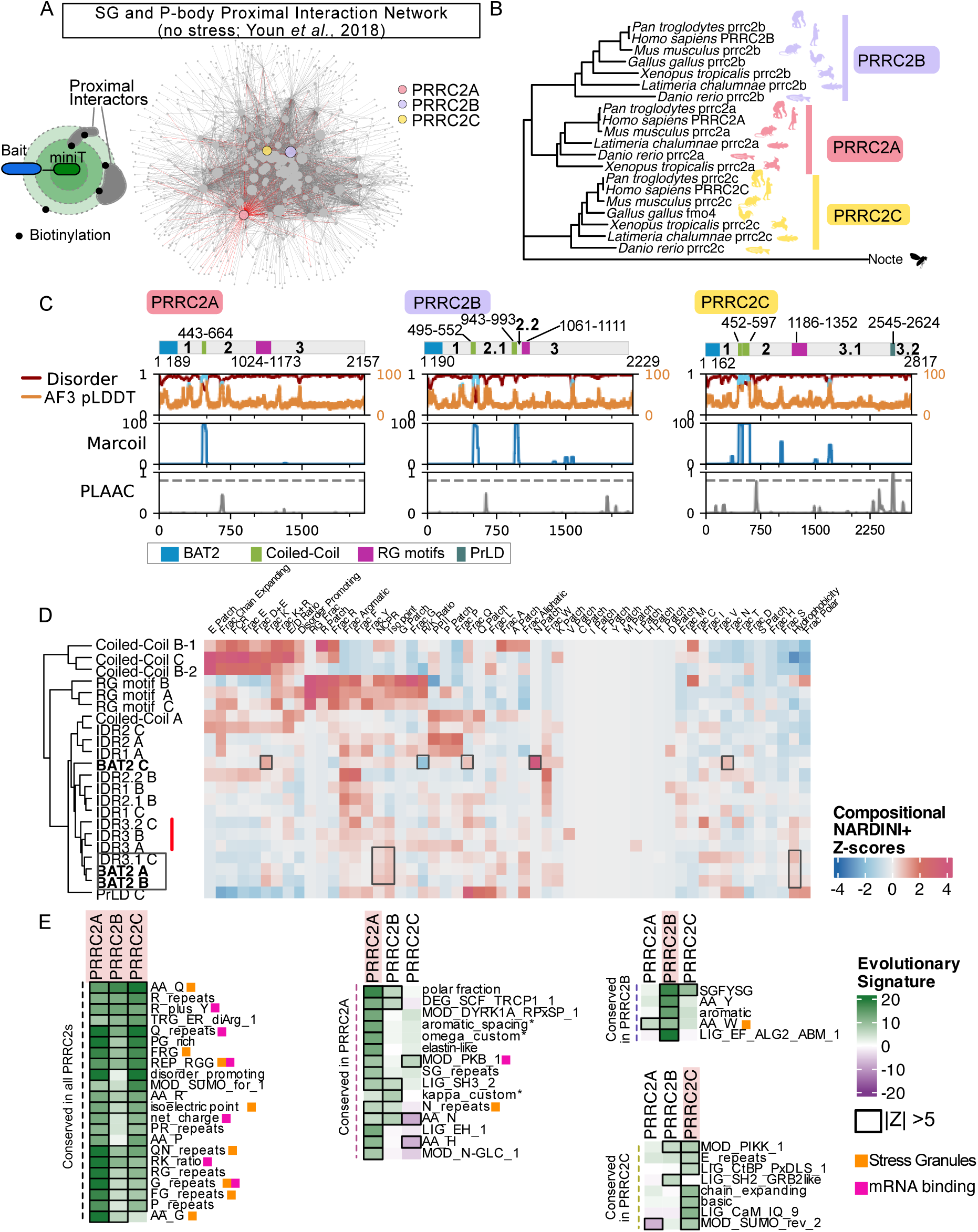
Molecular Features of Proline-Rich Coiled-Coils A (PRRC2A), PRRC2B and PRRC2C. **A)** Cytoscape network analysis identifies highly connected hub proteins within RNA granule-proximal interaction networks of cells under no stress, including PRRC2A, PRRC2B and PRRC2C. Proximal interactions between 59 RNA granule baits and 480 high-confidence interactors (AvgP≥0.95), reported in a previous study (Youn et al., 2018), are visualized as a network. Nodes represent RNA granule baits or high-confidence interactors, and edges denote proximal interactions. Node size scales with interaction degree, with highly connected proteins clustering toward the network center. PRRC2A (red), PRRC2B (blue), and PRRC2C (orange) show extensive connectivity to RNA granule baits (red edges). The schematic on the left illustrates how miniTurbo, fused to the bait protein of interest, biotinylates proximal interactors. **B)** Phylogenetic tree of PRRC2 orthologs. The rooting group is Nocte from *Drosophila melanogaster.* **C)** The domain architecture and disorder prediction analyses of the PRRC2 primary amino acid sequences. PRRC2 proteins share a conserved N-terminal domain referred to as ‘BAT2 domain’ (blue rectangle), putative coiled-coils (bright green rectangles), and RG motifs (magenta rectangle). PRRC2C contains a predicted Prion-Like Domain (dark rectangle). The top subplot shows Metapredict (V3.0) disorder prediction score in red, and the AlphaFold3 pLDDT score in orange. The pLDDT score indicates the per-residue prediction confidence. Regions with a pLDDT score > 70 (high confidence prediction) are coloured blue. The middle subplot shows the predicted coiled-coil probability from Marcoil. The bottom subplot shows the prion-like amino acid composition (PLAAC) score, with a dashed line indicating the significance threshold. **D)** The heatmap shows compositional Z-scores for IDR subregions within each PRRC2 protein, using the NARDINI+ analysis. Hierarchical clustering shows subregions with similar NARDINI+ Z-scores. The Z-scores indicate how much the molecular feature deviates from the mean of all human IDRs. **E)** Evolutionarily conserved molecular signature analyses of PRRC2 proteins. Conversed molecular features shared in all PRRC2 proteins and their orthologs (left), shared in PRRC2A orthologs (middle), in PRRC2B orthologs (right, top), and in PRRC2C orthologs (right, bottom). Conserved features with an absolute Z-score > 5 are highlighted with a border around the tile. Features predictive of stress granule localization are indicated with an orange box, and predictive of mRNA binding are indicated with a pink box next to each feature.

To identify proteins that mediate multivalent interactions important for SG formation, we selected interactors with the highest connectivity in the SG and P-body proximal interaction network. We extracted 59 BioID profiles of SG and P-body baits with high recovery of literature-curated RNA granule proteins (*i.e.,* ≥70% of high-confidence interactors with AvgP ≥ 0.95 being annotated as the consensus SG proteome, Tier 1^17,51^), which totalled 3,857 proximal interactions between 59 baits and 488 high-confidence preys. Network analysis with Cytoscape revealed high-confidence proximal interactors with a high degree of connectivity (**Fig. 1A**), including PRRC2 proteins, which each interacted with at least 42 out of the 59 baits (top 5%; **Fig. S1A**).

Evolutionary analyses revealed that PRRC2 proteins are conserved across jawed vertebrates (**Fig. 1B**). While we identified orthologs for all PRRC2 proteins across *Pan troglodytes* (chimpanzee), *Mus musculus* (mouse), *Xenopus tropicalis* (frog), and *Latimeria chalumnae* (coelacanth), we found no corresponding ortholog for PRRC2A in bird species like *Gallus gallus (*chicken). Unlike jawed vertebrates, which encode three paralogues of PRRC2 proteins, fruit flies, and other invertebrates encode only one ortholog (*e.g.* Nocte).

To infer function, we analyzed the domain architecture of PRRC2 proteins. All PRRC2 proteins and their orthologs share a BAT2 N-terminal domain (referred to as BAT2, InterPro family IPR009738), a ∼200-residue-long domain that lacks a stable structure with no known function (**Fig. 1C, S1B**). This domain is found in other eukaryotes^52^, such as *Trichoplax adhaerens* and *Arabidopsis thaliana.* In the middle of the protein, all PRRC2 proteins contain RG motifs. Using AlphaFold3 and Metapredict V3.0, we found that more than 90% of the primary sequences of PRRC2 proteins, spanning >2,000 residues, were predicted to be disordered (**Fig. 1C**). Furthermore, all PRRC2 orthologs exhibit high disorder, suggesting that disorder is an inherent feature of PRRC2 proteins and their orthologs (**Fig. S1B**).

The PRRC2 proteins also contain α helices that were confidently predicted by AlphaFold3 (pLDDT > 70; light blue in AF3 pLDDT plot **Fig. 1C**). Some of these α helices are predicted to have high propensity to form coiled-coils by Marcoil (**Fig. 1C**). Although predicted to form α helices, this region shows a high disorder score (>0.5 Metapredict score), suggesting that these α helices may fold conditionally. Prion-like amino acid composition (PLAAC) analysis identified regions with prion-like amino acid composition in all PRRC2 proteins; however, PRRC2C’s C-terminal region uniquely met the COREscore threshold (**Fig. 1B**). Analysis of PRRC2C orthologs revealed that the putative coiled-coils are conserved across PRRC2 orthologs (except for zebrafish Prrc2b), while the presence of prion-like regions above the stringent COREscore varied across the orthologs (**Fig. S1B**). Thus, PRRC2 paralogs are predicted to be intrinsically disordered proteins (IDPs) that possess coiled-coils and share similar domain architectures.

The lack of structured domains in PRRC2 proteins makes it difficult to predict their functions. However, various computational tools enabled us to analyze the properties of PRRC2 proteins based on their primary amino acid sequence, as described below.

#### 1) Conformational ensemble

IDPs adopt highly dynamic conformations and are best described as conformational ensembles. Characterizing the global conformational properties of IDPs can therefore provide biologically relevant insights into PRRC2 protein behavior. Parameters such as radius of gyration, end-to-end distance, asphericity and solvent accessible surface area inform whether an IDP adopts relatively expanded or compact conformations. Using AlphaFlex, a computational workflow that predicts atomistic conformational ensembles for proteins containing intrinsically disordered regions, we estimated these properties for the PRRC2 proteins^53^. The radius of gyration, end-to-end distance, and asphericity were highly similar among the PRRC2 proteins (**Fig. S1C**). Notably, the asphericity values were low for all PRRC2 proteins, suggesting a preference for relatively spherical rather than rod-like conformations. PRRC2C showed a slightly larger radius of gyration and end-to-end distance, consistent with its greater protein length, which scales with chain size^53^. Interestingly, PRRC2C exhibited substantially higher solvent-accessible surface area compared with PRRC2A and PRRC2B, suggesting it may be more accessible for interactions with other proteins and/or biomolecules.

#### 2) Molecular Grammar

Intrinsically disordered regions (IDRs) exhibit non-random composition and non-random patterning in their sequence, described as “molecular grammar”, which dictates their biophysical properties^54–58^. To examine how the grammar differs between paralogs, PRRC2 primary sequences were split into distinct regions using the boundaries of known domains and motifs; BAT2, IDR1, coiled-coil, IDR2, RG motif region, IDR3, PrLD (unique to PRRC2C) and IDR4 (unique to PRRC2C) (**Fig. 1C**). NARDINI+ calculates the compositional features of IDRs, and each feature is compared to the mean of all human IDRs (Z-score). By clustering the NARDINI+ compositional Z-scores for PRRC2 proteins, the similarities and differences between the PRRC2 proteins were revealed (**Fig. 1D; Table S1**).

Generally, the corresponding IDRs between the paralogs were similar and clustered together with some exceptions. The region containing RG motifs was the most similar among the paralogs, sharing similar features. Consistent with RG and RGG repeats’ ability to mediate interactions with RNA or proteins^59–62^, the RG motifs region in PRRC2A was shown to crosslink to RNA^33^. Sequence similarity in this RG motifs region suggest that PRRC2B and PRRC2C may interact with RNAs. In contrast to most corresponding regions clustering together, the BAT2 region of PRRC2C diverged from those of PRRC2A and PRRC2B and clustered with PRRC2B IDR1-2 instead. The BAT2 of the PRRC2A and PRRC2B clustered with the C-terminal IDR (IDR3.1) of PRRC2C. Thus, the BAT2 region of PRRC2C is predicted to exhibit properties different from those of the other paralogs.

#### 3) Evolutionarily conserved bulk molecular features

The ‘bulk’ molecular features of IDRs are conserved across evolution and are associated with function^55,56^. For example, analysis of the evolutionarily conserved bulk molecular features was performed on the full-length primary sequence of PRRC2 proteins, rather than on segmented regions, with minor modifications from the previous human IDRome analysis ^63^(see Methods). We expanded our evolutionary conservation analysis to ∼150 bulk molecular features, which include charge properties, physicochemical properties, repeats and short linear motifs (SLiMs). The Z-score represents the deviation from the expectation for disordered region evolution without constraints.

Fifty bulk properties were identified as significant in at least one of the three PRRC2 paralogs (|Z-score| >5, **Fig. 1E, Table S1**). Among the conserved signatures, 44% (22/50) of features were conserved in all three paralogs, including features commonly conserved in SG proteins^63^. Additionally, all PRRC2 proteins also showed high conservation in features associated with mRNA binding, suggesting all three paralogs are involved in mRNA binding and can localize to SGs. Each PRRC2 protein also has some molecular features that are uniquely conserved within their ortholog set (*e.g.* PRRC2C orthologs are conserved in a CtBP binding SLiM (LIG_CtBP_PxDL)). These sets of unique features suggest there is some functional divergence in the PRRC2 proteins.

Our computational analyses on the conformational ensembles, molecular grammar and features of the PRRC2 proteins highlight their structural and sequence similarities, as well as certain differences. Differences in their evolutionarily conserved molecular features may relate to the divergent function between paralogs, while similarities relate to their shared functions that are maintained, including features of SG proteins and mRNA binding.

### PRRC2 proteins localize to SGs with similar kinetics to the scaffold protein, G3BP1

Evolutionarily conserved feature analysis, together with the prior prey-centric BioID profiling performed in unstressed condition, predicted PRRC2 proteins as SG components. To validate this prediction experimentally, we examined PRRC2 proteins’ localization by immunofluorescence and live-cell imaging microscopy under two mechanistically distinct SG-inducing conditions: oxidative and hyperosmotic stress. Oxidative stress activates the ISR and induces the canonical SGs that depend on the scaffolding proteins, G3BP1 and G3BP2, whereas hyperosmotic stress is thought to promote SG assembly through molecular crowding and does not require G3BP1/2^64^. For immunofluorescence microscopy, endogenous PRRC2B and PRRC2C proteins were detected using antibodies that we validated for specificity. Because commercial antibodies with high-specificity were unavailable for PRRC2A, PRRC2A localization was assessed using inducibly and stably expressed 3×FLAG-tagged PRRC2A in PRRC2A-knockout cells to minimize overexpression.

Immunofluorescence microscopy revealed that, like G3BP1—which exhibits biphasic organization with a stable, dense “core” and a more labile peripheral “shell”—PRRC2 proteins showed high-intensity subregions within SGs, analogous to the “cores” (**Fig. 2A**)^65^. All three PRRC2 proteins robustly colocalized with the SG marker G3BP1 under both oxidative and hyperosmotic stress conditions (**Fig. 2A**). Quantitative analysis using the Manders’ overlap coefficients demonstrated near-complete co-occurrence between each PRRC2 protein and G3BP1 signals in both stress paradigms (mean Manders’ coefficients > 0.9; **Fig. S2A**), with reciprocal overlap comparable to that observed for G3BP1 with itself or with UBAP2L, a well-characterized G3BP1 interactor and SG protein. Together, these data indicate that the PRRC2 proteins spatially coincide with G3BP1 across distinct modes of SG assembly.

**Figure 2.**
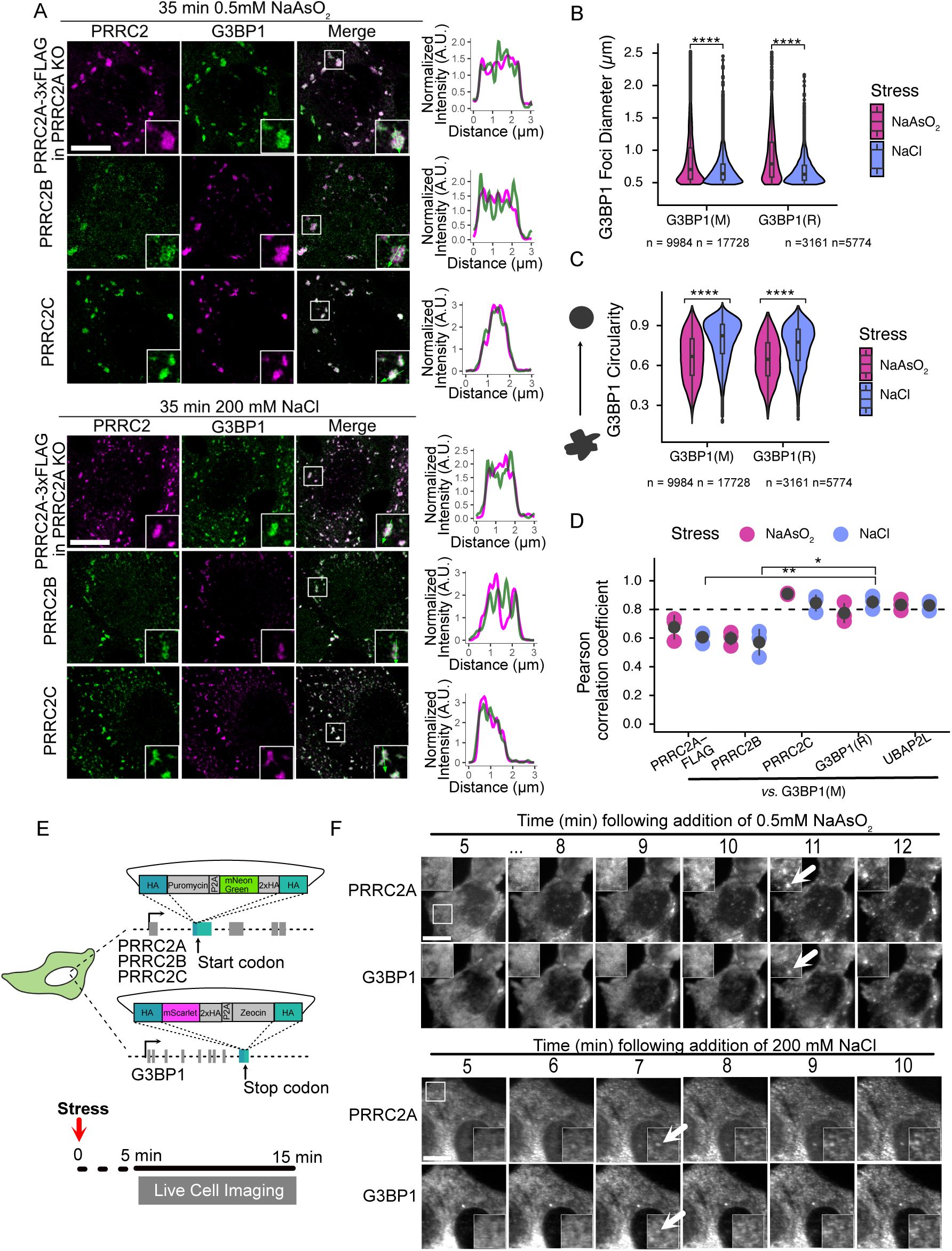
Stress-induced localization of PRRC2 proteins to SGs and their condensation kinetics. **A)** Representative immunofluorescence max intensity projections show PRRC2A-3xFLAG, endogenous PRRC2B and endogenous PRRC2C localization upon 35 min of oxidative (0.5 mM NaAsO_2_; top) and hyperosmotic (200 mM NaCl; bottom) stress in HeLa Flp-In T-REx cell lines. The PRRC2A-3xFLAG was inducibly expressed in PRRC2A knockout (KO) cells to minimize overexpression. Right: The line-scan plots show the signal intensities of each PRRC2 protein and G3BP1 in the selected stress granule, as shown in the insets. Scale bar: 20 µm. **B)** The violin plot shows the distribution of G3BP1 foci diameter (µm) under oxidative (NaAsO_2_) or hyperosmotic (NaCl) stress, captured using two different antibodies; letters in parentheses denote the G3BP1 monoclonal mouse antibody (M) or polyclonal rabbit antibody (R). Wilcoxon rank-sum test comparing G3BP1 foci diameter between SGs induced by the two stressors. **** *P* <0.0001. Quantified from three biological replicates. n = number of granules quantified. **C)**The violin plot shows the degree of circularity for G3BP1 foci formed under oxidative (NaAsO_2_) or hyperosmotic (NaCl) stress, captured using two different antibodies; letters in parentheses denote the G3BP1 monoclonal mouse antibody (M) or the polyclonal rabbit antibody (R). Wilcoxon rank-sum test comparing G3BP1 foci diameter between SGs induced by the two stressors. **** *P* <0.0001. Quantified from three biological replicates. n = number of granules quantified. **D)** The plot shows the Pearson correlation coefficient (PCC) analysis of fluorescence intensity between the indicated proteins. PCC was calculated across the entire image. The black circle and line represent the mean and standard deviation, respectively. Statistics were performed using Student’s t-test with Holm adjustment, * *P* < 0.05, ** *P* < 0.01. Data from three biological replicates. The dashed line marks PCC = 0.80. n = number of granules quantified. **E)** The top schematic illustrates the CRISPR-Cas9-mediated homology-directed repair strategy employed to generate endogenously fluorescence protein-tagged HAP1 cells. PRRC2 proteins were endogenously tagged with mNeon Green at the N-terminal. G3BP1 was tagged at the C-terminal end with mScarlet. The bottom schematic shows the duration of live-cell imaging, beginning 5 minutes after the addition of a stressor. Cells were imaged for 10 minutes with 30-second intervals using Lattice light-sheet microscopy. **F)** Representative max intensity projections of HAP1 cells endogenously expressing mNeonGreen-PRRC2A and G3BP1-mScarlet. The white arrow in the inset points to newly formed SGs. Scale bar: 10 µm.

Stress conditions differentially affected the size and morphology of SGs. Granules induced by oxidative stress were generally larger (*e.g.,* median diameter of 0.79 mm *vs* 0.63 mm, G3BP1(R)) and less circular (*e.g.,* median circularity 0.65 *vs* 0.78, G3BP1(R)) than those formed under hyperosmotic stress (**Fig. 2B-C**). PRRC2-positive signal exhibited a similar pattern, with comparable changes in diameter and circularity across stress conditions **(Fig. S2B)**. These observations suggest that distinct intracellular environments associated with different stress modalities modulate the architecture of SGs.

Line scan analysis comparing PRRC2 protein intensity profiles to G3BP1 revealed paralog-specific differences in their spatial distributions within SGs. Under both oxidative and hyperosmotic stress, the maximal intensity peaks of PRRC2C closely coincided with those of G3BP1, whereas PRRC2A or PRRC2B showed less concordance, particularly under hyperosmotic stress, where their peak intensities were visibly offset from G3BP1 (**Fig. 2A**). Consistent with these observations, quantitative assessment using Pearson’s correlation coefficient (PCC) across entire images revealed a strong correlation between PRRC2C and G3BP1 signals in both stress conditions (mean PCC > 0.8), comparable to the correlation of G3BP1 with itself or with UBAP2L (**Fig. 2D**). In contrast, PRRC2A and PRRC2B exhibited more modest correlations with G3BP1 in both stress contexts (mean PCC ≈0.6 for both), with significantly lower PCC values than G3BP1 self-correlation. As for PRRC2B, this can be attributed to its subpopulation localizing to the nucleus, which we discuss further in the next section. Together, these analyses indicate that PRRC2C preferentially occupies the same SG subcompartment as G3BP1, whereas PRRC2A and PRRC2B are spatially enriched in distinct subregions within SGs.

To investigate the assembly kinetics of PRRC2 proteins in SGs, we performed live cell imaging analysis of PRRC2 paralogs and G3BP1 following stress induction. Endogenously tagged PRRC2 proteins (mNeonGreen) and G3BP1 (mScarlet) expressed in near-haploid, HAP1 cells were imaged using lattice light-sheet microscopy (**Fig. 2E-F, S2C**). Under oxidative stress, PRRC2A condensation was detected approximately 10-11 min after stress induction, while under hyperosmotic stress, condensation occurred at ∼6-7 min; in both cases, PRRC2A recruitment coincided with the onset of SG formation as marked by G3BP1 condensation (**Fig. 2F; Video S1-2**). PRRC2B and PRRC2C displayed similar assembly kinetics under both stress conditions (**Fig. S2C, Video S3-6**). Slight delay in estimated timing of G3BP1 condensation observed here following oxidative stress relative to previous reports^66^ (∼5 min using lattice light-sheet microscopy) can be attributed to differences in G3BP1 expression strategy (endogenous tagging *vs.* BAC-based stable expression) or cell type (*i.e.,* HAP1 *vs.* HeLa). Together, these data demonstrate that all three PRRC2 paralogs are recruited to SGs at the earliest detectable stages of SG assembly.

### Functional proteomics analysis reveals shared interactions of PRRC2 proteins with translation machinery

Protein-protein interaction networks provide insight into protein function through guilt-by-association with its interacting partners. Previous studies have applied functional proteomics approaches to PRRC2A or PRRC2B; however, differences in experimental strategies and cellular contexts limit direct comparison across datasets PRRC2 proteins ^31,32,67^.

To define systematically shared and paralog-specific molecular functions of the PRRC2 protein family, we employed affinity purification coupled with mass spectrometry (AP-MS). AP-MS captures stable protein-protein interactors that withstand the affinity purification, including both direct and indirect interactions. AP-MS experiments were performed in HEK293 Flp-In T-REx cells stably expressing inducible 3×FLAG-tagged PRRC2 proteins (**Fig. 3A**). Samples were analyzed by Data-Independent Acquisition (DIA) mass spectrometry and processed using DIA-NN to improve protein identification and quantitative accuracy^68^ (see Methods). Biological replicates showed high reproducibility (CV<15%, **Table S2**). High-confidence interactors were defined using SAINTq^69^ by performing statistical analysis against parallel negative controls acquired (**Fig. 3A**; Methods). High-confidence AP-MS interactors were defined using a threshold of average SAINT score (AvgP) ≥0.98, based on our prior analyses of SG proteins^17^.

**Figure 3.**
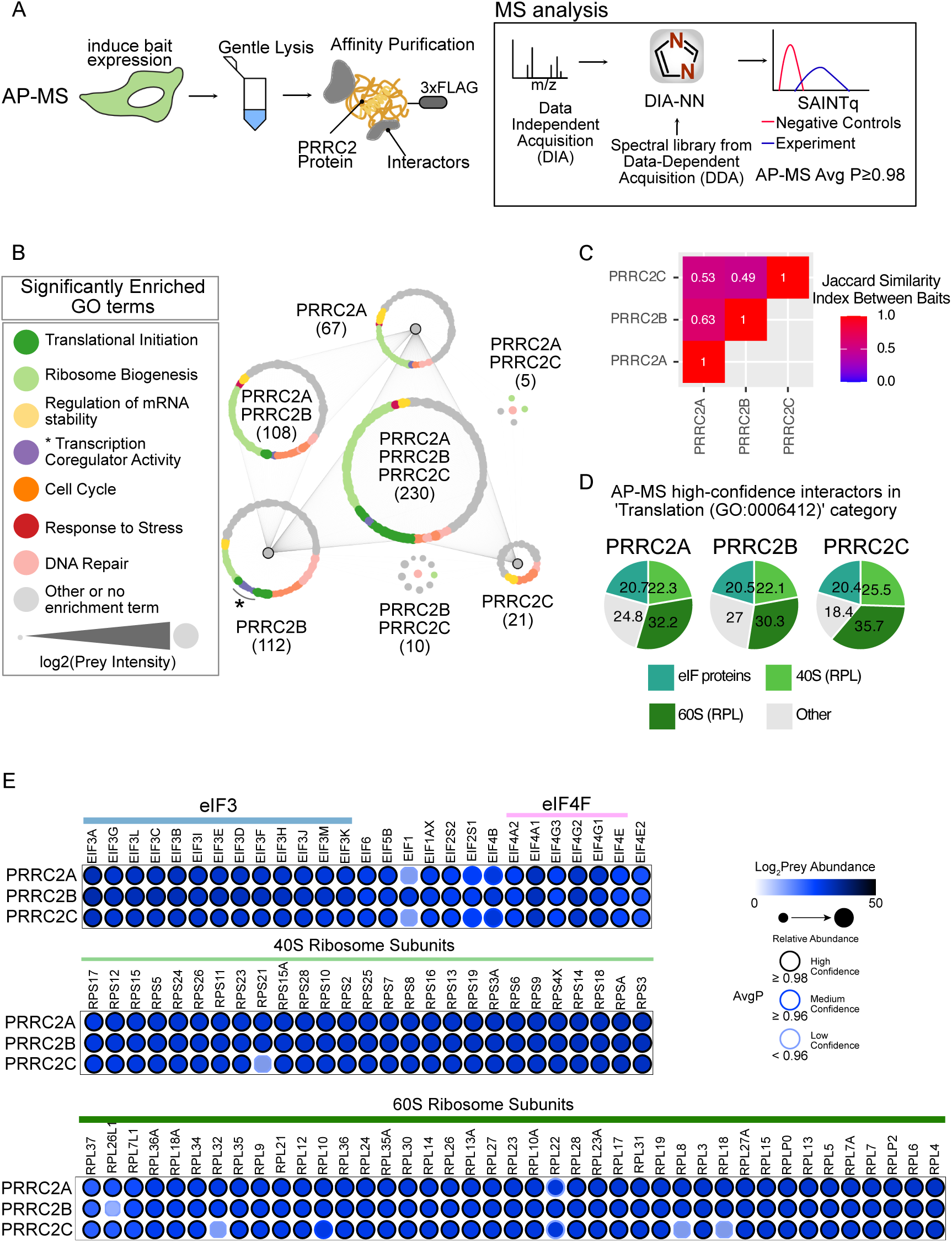
AP-MS Reveals Shared Interactions with Ribosomes and Translation Initiation Factors. **A)** Schematics display the experimental workflow for AP-MS. Samples were analyzed in data-independent acquisition mode on the mass spectrometer, which is processed using DIA-NN, and analyzed using SAINTq to determine high-confidence interactors (right panel). **B)** Cytoscape network visualization displays high-confidence proximal interactors (prey) identified by PRRC2A, PRRC2B, and PRRC2C using AP-MS. Coloured nodes represent interactors assigned to a significantly enriched gene ontology (GO) term, with node colour indicating the corresponding GO category. Lines between nodes represent a high-confidence protein-protein interaction. Node size reflects prey abundance (log_2_ (prey intensity)). The number of interactors unique to each paralog or shared among PRRC2A, PRRC2B, and PRRC2C is indicated in brackets. **C)** The heatmap shows the Jaccard similarity indices of high-confidence interactors identified in AP-MS between the PRRC2 baits. **D)** The pie chart shows the composition of high-confidence interactors in the ‘GO:006412 Translation’ category in AP-MS datasets. **E)** The dot plots show PRRC2 protein interactors involved in translation initiation or components of ribosomal subunits.

AP-MS profiling revealed a high degree of similarity among PRRC2 paralogs. PRRC2A, PRRC2B, and PRRC2C identified 403, 460, and 266 high-confidence stable interactors, respectively, with substantial overlap across datasets (**Fig. 3B, Table S2**). While all paralogs share interactions with components involved in translation and ribosome biogenesis, PRRC2B differed from the other two paralogs, displaying greater associations with proteins harboring ‘transcription coregulator activity’ (**Table S3**). Pairwise Jaccard indices of the PRRC2 proteins’ AP-MS profiles ranged from 0.49 to 0.63, highlighting their subtle differences (**Fig. 3C**).

Interactors shared among all three PRRC2 proteins were enriched for factors involved in translation initiation and ribosome biogenesis (**Fig. 3B**), implicating conserved roles for PRRC2 proteins in these processes. Amongst high-confidence AP-MS interactors annotated under the Gene Ontology term, ‘Translation (GO:0006412)’, the majority corresponded to core components of the translational apparatus (**Fig. 3D**). Furthermore, all three PRRC2 proteins interacted with the eIF4F complex, the eIF3 complex, 26 small ribosomal subunits, and 39 large ribosomal subunits (**Fig. 3E**). Notably, translation elongation and translation termination factors were not detected, suggesting that PRRC2 proteins are specifically participating in translation initiation rather than all stages of translation.

### Dynamic Interaction Networks of PRRC2A and PRRC2B During Stress Reveal Their Context-Dependent Functions

Microscopic formation of SG is associated with dynamic rearrangements in the protein interaction networks of SG proteins^70^. PRRC2 proteins exhibit extensive connectivity with SG proteins under basal conditions (Fig. 1A). To determine how the interaction networks of PRRC2 proteins are rearranged upon stress, proximity labeling experiments of PRRC2C proteins were performed with cells under no stress, 5 min into oxidative stress (0.5 mM NaAsO₂) or 5 min into hyperosmotic stress (200 mM NaCl) for 30 min (**Fig. 4A**). BioID was performed in HEK293 Flp-In T-REx cells stably expressing inducible miniTurbo-tagged and the mass spectrometry data was analyzed in DIA-NN and SAINTq (**Fig. 3A**). Biological replicates showed high reproducibility (**Table S4**), and an AvgP ≥0.95 threshold was used to define high-confidence proximal interactors.

**Figure 4.**
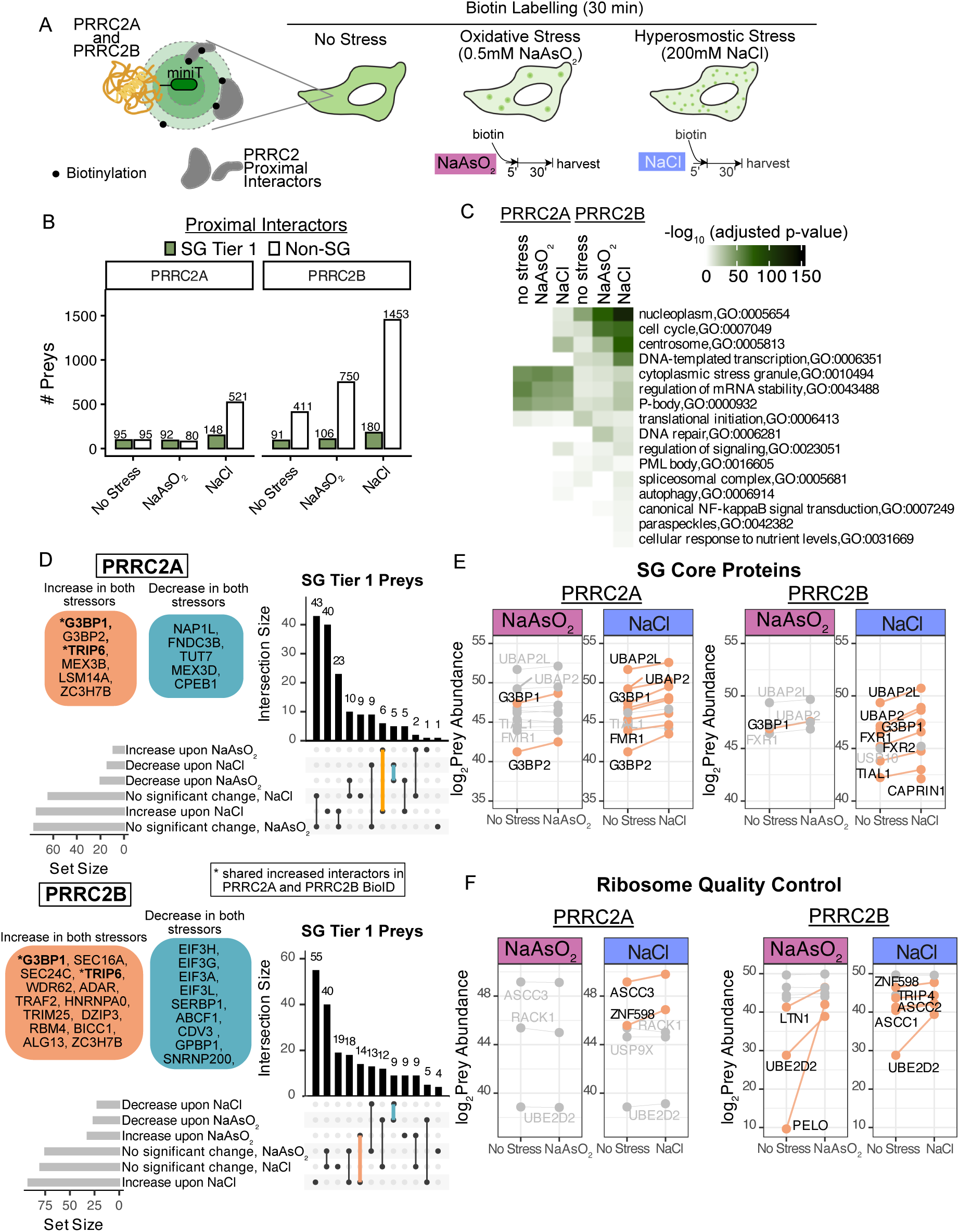
Stress Induces Dynamic Rearrangement in BioID Profiles of PRRC2A and PRRC2B. **A)** The schematic shows BioID labelling during three conditions: no stress, oxidative stress and hyperosmotic stress. For stress BioID, biotin was added 5 minutes into stress with either 0.5 mM NaAsO_2_ or NaCl to label proximal interactors during SG formation. MS analysis is identical to AP-MS. AvgP ≥ 0.95 was used as a cutoff for high-confidence proximal interactors. **B)** The bar plots show the number of high-confidence interactors identified by PRRC2A or PRRC2B in no stress, oxidative stress, and hyperosmotic stress conditions. Identified interactors are divided into SG tier 1 proteins (green) and non-SG tier 1 proteins (white). **C)**Heatmap visualizes the statistical significance of GO terms associated with PRRC2A and PRRC2B proximal interactors in each indicated condition. **D)** The upset plots show the intersections of high-confidence interactors that increase, decrease, or remain unchanged upon the indicated stress in PRRC2A or PRRC2B BioID experiments (fold change ≥|1.5-fold|). Preys showing differential proximal interactions in both stress conditions are listed: increased group in the orange box and decreased group in the purple box. Preys identified by both PRRC2A and PRRC2B and show changes in the same direction are in bold with *. **E)** The line plot shows changes in abundance for well-characterized SG proteins identified as high-confidence interactors in either no-stress or stress conditions. Preys whose abundance changes by ≥ 1.5-fold are indicated in black with orange lines, and those below this threshold are indicated in gray. Changes in abundance indicate changes in proximial interactions. **F)** The line plot shows the abundance changes for proteins involved in ribosome quality control, identified as high-confidence interactors in either the no-stress or stress conditions. Preys whose abundance changes by ≥ 1.5-fold are indicated in black with orange lines, and those below this threshold are indicated in gray. Changes in abundance indicate changes in proximal interactions.

Under no stress, BioID profiling identified 184, 500, and 14 high-confidence proximal interactors (*i.e.,* preys) for PRRC2A, PRRC2B, and PRRC2C, respectively (**Table S4**). Subsequent analyses focused on PRRC2A and PRRC2B, as PRRC2C BioID yielded limited information due to low protein expression and inefficient biotinylation despite testing multiple conditions (**Fig. S3A,** see Methods). PRRC2A and PRRC2B BioID datasets shared only a small subset of proximal interactors with each other, with a Jaccard similarity index of 0.11 (scale: 0 to 1; **Fig. S3B**). The overlap between AP-MS and BioID datasets was limited (Jaccard indices 0.03-0.08) due to the distinct interaction modalities captured by each method, consistent with previous reports^48,71^ (**Fig. S3B**). However, both approaches revealed strong associations between PRRC2 proteins and the translation machinery under basal conditions (**Table S3, S5**).

Immunofluorescence microscopy confirmed that BioID labeling of PRRC2A and PRRC2B during stress resulted in strong enrichment of biotinylation signal in SGs (**Fig. S3C**). Proteomics analysis showed that stress context had a major influence on their proximal interaction networks. Hyperosmotic stress caused a substantial increase in the number of high-confidence interactors for both PRRC2A and PRRC2B, mostly due to novel interactions with non-SG proteins (**Fig. 4B**). This observation is consistent with what was observed in stress BioID data for other SG proteins^70^. Overall, the increase in the number of proximal interaction partners identified by PRRC2B during stress was greater than that of PRRC2A; we identified a total of 518 and 1,416 novel stress-induced proximal interactions for PRRC2A and PRRC2B stress BioID, respectively. About 50 % of literature curated SG proteome (Tier 1) interactors identified by PRRC2A were shared with those identified by PRRC2B, indicating PRRC2A and PRRC2B share some redundancy in the SG network (**Fig. S4A**).

GO enrichment analysis of high-confidence interactors identified by PRRC2A and PRRC2B in each condition summarizes the changes in their BioID profiles, highlighting the functional divergence of PRRC2B from PRRC2A in cellular response to stress (**Fig. 4C, Table S5**). Heatmap visualization of significantly enriched GO terms demonstrates that the PRRC2A BioID profile remains largely the same across all conditions, with new interactions during hyperosmotic stress. PRRC2A is predominantly associated with SG proteins and proteins involved in the regulation of mRNA stability and translation initiation regardless of the cell state, except for novel proximal interactors found under hyperosmotic stress (*e.g.,* proteins involved in the regulation of signalling, centrosome and cell cycle). In contrast, the PRRC2B BioID profile shows considerable stress-induced changes. Like PRRC2A, PRRC2B shows a stronger association with proteins that localize to the centrosome in hyperosmotic stress. PRRC2B shows enhanced engagement with nucleoplasm and cell cycle proteins in both conditions, while PRRC2B uniquely associates with DNA repair proteins upon oxidative stress. In summary, BioID profiling during stress revealed functional differences between PRRC2A and PRRC2B and demonstrated the PRRC2B’s versatility in function during stress adaptation.

Next, we examined the dynamic interaction changes with SG components. Consistent with earlier observations (**Fig. 1A**), an extensive network of protein-protein interactions was detected between PRRC2A and PRRC2B proteins and SG proteins; each bait identified ∼20% of the literature-curated SG proteome (Tier 1, 407 total) in BioID basal condition (**Fig. 4B**). Examination of dynamic proximal interactions between SG tier 1 proteins and PRRC2 proteins revealed that the majority of interactions remain largely unchanged, consistent with other reports^17,18,70^ (**Fig. 4D**). However, PRRC2A and PRRC2B both showed increased proximal interactions with G3BP1, the SG scaffold, regardless of the stress imposed. Given G3BP1’s role in creating multivalent interactions that facilitate SG formation^13,72^, this suggests that PRRC2A and PRRC2B become engaged in G3BP1’s interaction network during SG formation.

To assess whether PRRC2A and PRRC2B exhibit increased proximal associations with G3BP1-interacting proteins—CAPRIN1, UBAP2L, and USP10—as well as other established SG proteins (TIAL1, UBAP2, FMR1, FXR1 and FXR2), their abundances between unstressed and each stressed condition were analyzed. Under oxidative stress, G3BP1/G3BP2 were the only interactors showing a ≥1.5-fold increase, while in hyperosmotic stress, multiple SG proteins displayed ≥1.5-fold increase (**Fig. 4E**). These results indicate that BioID profiling of PRRC2A and PRRC2B captures context-dependent remodelling of proximal interaction networks during SG assembly.

Given the association of PRRC2 proteins with translational machinery, stress-induced changes in proximal interactions between PRRC2 proteins and translation-related components were examined. Under both oxidative and hyperosmotic stress, PRRC2B showed dynamic proximal interactions with subunits of the eIF3 complex (**Fig. S4B**). Oxidative stress induced mixed changes; proximal interactions with eIF3e, eIF3k and eIF3i increased, while interactions with eIF3a, eIF3g, eIF3l, eIF3f, eIF3h, and eIF3j were reduced. During hyperosmotic stress, most subunits showed decreased proximal interactions with PRRC2B. Given eIF3 complex’s known role in selective mRNA translation during stress^42,43,73,74^, this differential interaction patterns may reflect spatial re-arrangement between PRRC2B and the eIF3 complex during stress-induced translation reprograming.

Proximal interactions between PRRC2A or PRRC2B with additional translation initiation factors and ribosomal subunits were examined. PRRC2A and PRRC2B displayed distinct, stress-specific interaction profiles (**Fig. S4C-D**). PRRC2A showed reduced associations with eIF4G proteins and ribosomal subunits under oxidative stress, but increased associations under hyperosmotic stress. In contrast, PRRC2B displayed decreased interactions with eIF4G1 during oxidative stress and with eIF4G2 during hyperosmotic stress. Proximal interactions between PRRC2B and ribosomal subunits were largely preserved across stress conditions, although individual subunits displayed stress-dependent increases or decreases in association. Together, these patterns suggest that PRRC2A and PRRC2B differentially engage with translation machinery in a stress-dependent manner.

In addition to translation initiation factors, both PRRC2A and PRRC2B showed pre-existing proximal interactions with components of the ribosome quality control pathway, which increased upon both oxidative and hyperosmotic stress conditions (**Fig. 4F**). Cellular stress can damage RNA, which slows translation, and promotes ribosome collisions, thereby activating the ribosome quality control pathway^75,76^. In this pathway, stalled ribosomes are ubiquitinated by ZNF598 and split by the ASC-1 complex (ASCC1/2/3 and TRIP4). The nascent polypeptide chain is targeted for proteasome-mediated degradation by LTN1, while PELO facilitates ribosome recycling. PRRC2A showed increased proximal associations with ASCC3 and ZNF598 specifically during hyperosmotic stress, whereas PRRC2B showed enhanced interactions with multiple ribosome quality control components under both stress conditions.

Concomitant with activation of ribosome quality control, the damaged mRNAs are subsequently cleared through the canonical RNA decay pathway, involving mRNA decapping, which involves DDX6, DCP1A, DCP1B, EDC3 and EDC4, followed by 5’—3’ degradation by the exoribonuclease, XRN1. Both PRRC2A and PRRC2B displayed pre-existing interactions with these decay factors, which increased specifically during hyperosmotic stress (**Fig. S4E**). These observations suggest that PRRC2A and PRRC2B are associated with components in ribosome quality control and mRNA decay pathways in response to stress.

Overall, our results reveal dynamic, stress-dependent remodelling of PRRC2A-and PRRC2B-associated proximal interaction networks. The dynamic changes observed upon stress support a role for PRRC2 proteins in stress-induced translation reprogramming, ribosome quality control and mRNA surveillance.

### Microscopy and functional proteomics reveal PRRC2B’s dual localization to paraspeckle with potential functions in DNA damage and the cell cycle

Coordination of biological processes in the nucleus and cytoplasm is important for the regulation of gene expression, and this regulation can be mediated by the coordination of nuclear and cytoplasmic condensates. Supporting this idea, many components of the cytoplasmic stress granules show dual localization to nuclear ribonucleoprotein complexes (Youn et al., 2018), and a SG protein, UBAP2L, has been shown to influence nuclear condensates under oxidative stress^77^. Hence, we further sought to assess PRRC2B’s divergent function identified in its AP-MS and BioID profiles and whether its nuclear localization induced changes in protein interactions.

Stress expanded the number of interactors associated with PRRC2B, which were enriched for proteins residing in the nucleoplasm (**Fig. 4C**). Concomitantly, stress induced an increase in the number of associations between PRRC2B and various nuclear importers in both stress conditions (**Fig. S5A**), suggesting that PRRC2B translocate to the nucleus. Indeed, putative nuclear localization signals (NLS) and putative PY-nuclear localization signals (PY-NLS) were found in the PRRC2 proteins, with PRRC2B containing two classical NLS and one PY-NLS—the highest number of putative NLS sequences out of the three paralogs (**Fig. S5B**), which likely contributes to PRRC2B’s higher degree of nuclear localization compared to other paralogs^78^.

Consistent with the BioID results, immunofluorescence microscopy experiments showed PRRC2B’s nuclear puncta in a subset population of cells under non-stress conditions, which is morphologically altered under oxidative stress (**Figure S5C**). Next, we validated whether these increased nuclear protein interactions occur in the nucleus. Three nuclear interactors showing increased association during oxidative stress were selected (BEND3, BCOR and eIF3e) (**Fig. S5D**). Colocalization was assessed using immunofluorescence microscopy of HEK293 cells stably expressing 3×FLAG-tagged PRRC2B in inducible manner and transiently expressing fluorescently tagged interactors in untreated and stressed cells (0.5mM NaAsO_2_). Under no stress, PRRC2B forms nuclear puncta in some cells, while the nuclear interactors generally remain diffuse (**Fig. 5A**). During oxidative stress, PRRC2B localized to large stress granules along with nuclear punctate structures. All three preys form puncta; BCOR and BEND3 formed puncta in the nucleus, whereas eIF3e formed large puncta in the cytoplasm and small puncta in the nucleus, like PRRC2B. BCOR showed high degree of overlap in signal with PRRC2B, while BEND3 and eIF3e puncta were often juxtaposed to PRRC2B. This confirms PRRC2B’s increased interactions with BEND3, BCOR and eIF3e occur in the nucleus and this is correlated with their localization to punctate structures.

**Figure 5.**
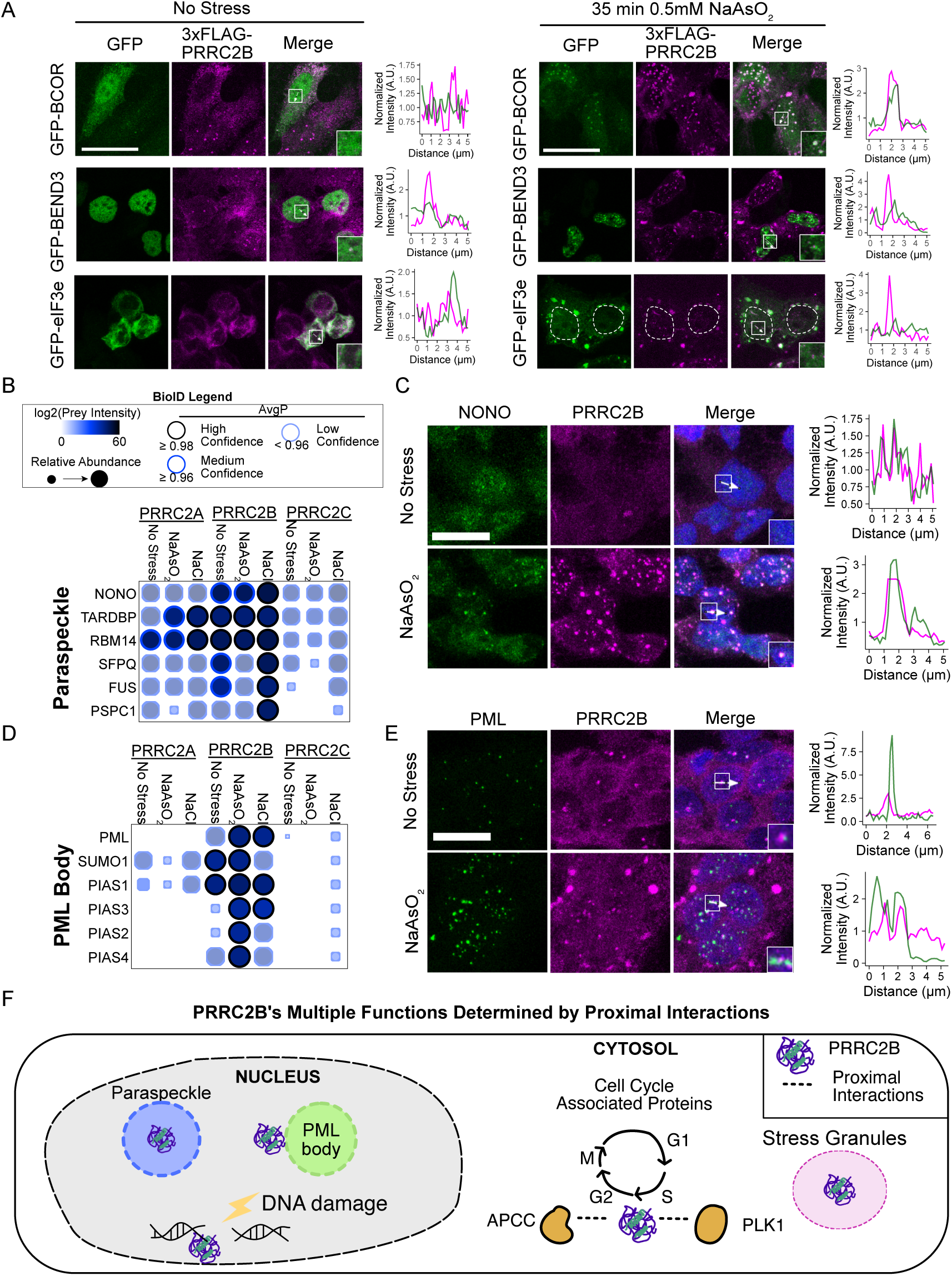
PRRC2B’s Divergent Functions Predicted from Microscopy and Functional Proteomics. **A)** Representative immunofluorescence max intensity projections show transiently transfected GFP-tagged nuclear preys and 3x-FLAG-tagged PRRC2B puncta in HEK293 Flp-In T-REx cell lines. The white arrow indicates the line used to draw the intensity profile. Scale bar: 20 µm. The inset shows the region or foci where the line was drawn (white box). The white dashed line indicates the boundary for the nucleus in the GFP-eIF3e immunofluorescence images. **B)** The dot plot shows high-confidence prey known to localize to paraspeckles. **C)** Representative immunofluorescence max intensity projections show the localization of NONO and PRRC2B in HEK293 Flp-In TREx cells stably expressing 3xFLAG-tagged PRRC2B, as detected with antibodies targeting each protein (anti-NONO and anti-PRRC2B). The white arrow indicates the line drawn for the intensity profile. Scale bar: 10 µm. The inset shows the region or foci where the line was drawn (white box). **D)** The dot plot shows PML, SUMO and E3 SUMO ligases. **E)** Representative immunofluorescence max intensity projections show the localization of PML and PRRC2B in HEK293 Flp-In TREx cells stably expressing 3xFLAG-tagged PRRC2B, as detected with antibodies targeting each protein (anti-PML and anti-PRRC2B). The white arrow indicates the line used to draw the intensity profile. Scale bar: 10 µm. The inset shows the region or foci where the line was drawn. **F)** PRRC2B’s multiple functions are determined by proximal interactions. PRRC2B is dually localized to the nucleus. Stress induces increased proximal interactions with nuclear proteins. PRRC2B can localize into paraspeckles and dock next to PML bodies. During oxidative stress, PRRC2B shows increased proximal interaction with DNA damage proteins. In the cytosol, PRRC2B localizes to SGs during stress. Stress also induces increased proximal interactions with anaphase-promoting complex (APCC) and polo-like kinase 1 (PLK1).

PRRC2B also showed proximal interactions with components of nuclear condensates, specifically, paraspeckles and PML bodies. Proximal interactions with paraspeckle and PML body components were unique to PRRC2B, which either sustained or increased upon stress (Fig. **5B-C**). To test PRRC2B’s colocalization with these condensates, we examined their signal overlap under oxidative stress, which is known to induce increased assembly of paraspeckles^77^ and PML bodies^79^. Under no stress, PRRC2B colocalization with the paraspeckle marker, NONO, was more diffuse and difficult to assess. However, upon oxidative stress, PRRC2B displayed clear colocalization with NONO (**Fig. 5D**). Consistently, BioID profiling-based analysis of the nuclear compartments also predicted PRRC2B as a paraspeckle protein^80^. This confirms that PRRC2B exhibits dual localization to nuclear paraspeckles and cytoplasmic stress granules.

While distinct from paraspeckles, PML bodies, containing PML protein and various sumoylated proteins, are involved in responses to DNA damage and apoptotic signal^81^. PRRC2B’s proximal interactions with the PML body components (PML, SUMO1, PIAS1-4) become most pronounced under oxidative stress (**Fig. 5D**). Under no stress, PRRC2B docks next to PML bodies, which remains the same as the number of PML bodies increases upon oxidative stress (**Fig. 5E**). Given that oxidation of the scaffold protein, PML, increases sumoylation (SUMO1) and SUMO-SIM interactions^79^, PRRC2B’s increase in association with these factors likely reflect the structural changes occurring with PLM bodies and these interactions are due to PRRC2B’s spatial proximity to PLM bodies.

Oxidative stress induces DNA damage^82^. Consistent with PRRC2B’s proximity to PML bodies, which are found at the sites of DNA damage^83^, PRRC2B uniquely displayed proximal interactions with proteins involved in DNA damage response and repair, which were enhanced upon oxidative stress, including TP53, RAD451C and H2AX (**Fig. S5E-F**).

PRRC2B has been implicated in cell cycle functions; it promotes translation of cell cycle proteins^31^, and itself is shown to be rapidly degraded, suggesting that it can be regulated by the cell cycle^84^. Supporting this, our BioID analysis showed PRRC2B association with cell cycle regulators, including polo-like kinases 1 (PLK1), M-phase inducer phosphatase 3 (CDC25C) and anaphase-promoting complex proteins (APCC) (**Fig. S5G-H**). Furthermore, these interactions were enhanced in both stress conditions, including novel interactions with a subset of proteasome subunits, suggesting PRRC2B’s turnover during cellular stress response. Supporting PRRC2B’s interaction with APCC and high protein turnover rate, a degron sequence, DEG_APCC_KENBOX2 (Anaphase Promoting Complex destruction sequence), was identified in PRRC2B (**Fig. S5I**). While the DEG_APCC_KENBOX2 sequence is also found in PRRC2C, this SLiM is only strongly conserved in PRRC2B orthologs.

Distinct from its paralogs, these data support that PRRC2B has additional divergent roles (**Fig. 5F**). Its proximal interactome uniquely includes nuclear transcriptional regulators, DNA damage response factors, and nuclear condensate components such as paraspeckles and PML bodies, with stress-dependent expansion of these interactions. Together with prior evidence linking PRRC2B to alternative splicing under hypoxia^36^, these findings suggest that PRRC2B coordinates nuclear RNA regulation with cytoplasmic translation control.

### PRRC proteins contribute to SG assembly in a context-specific manner

PRRC2C was shown to promote SG assembly under oxidative stress^17^. To determine if other PRRC2 paralogs redundantly promote SG assembly, we generated CRISPR Cas9-mediated knockout clones for single, double or triple mutants sequentially in HeLa Flp-In TREx cells (**Fig. 6A**). Single clones were isolated, and successful frameshift mutants arising from erroneous repair were validated by genotyping (**Table S6;** see Methods).

**Figure 6.**
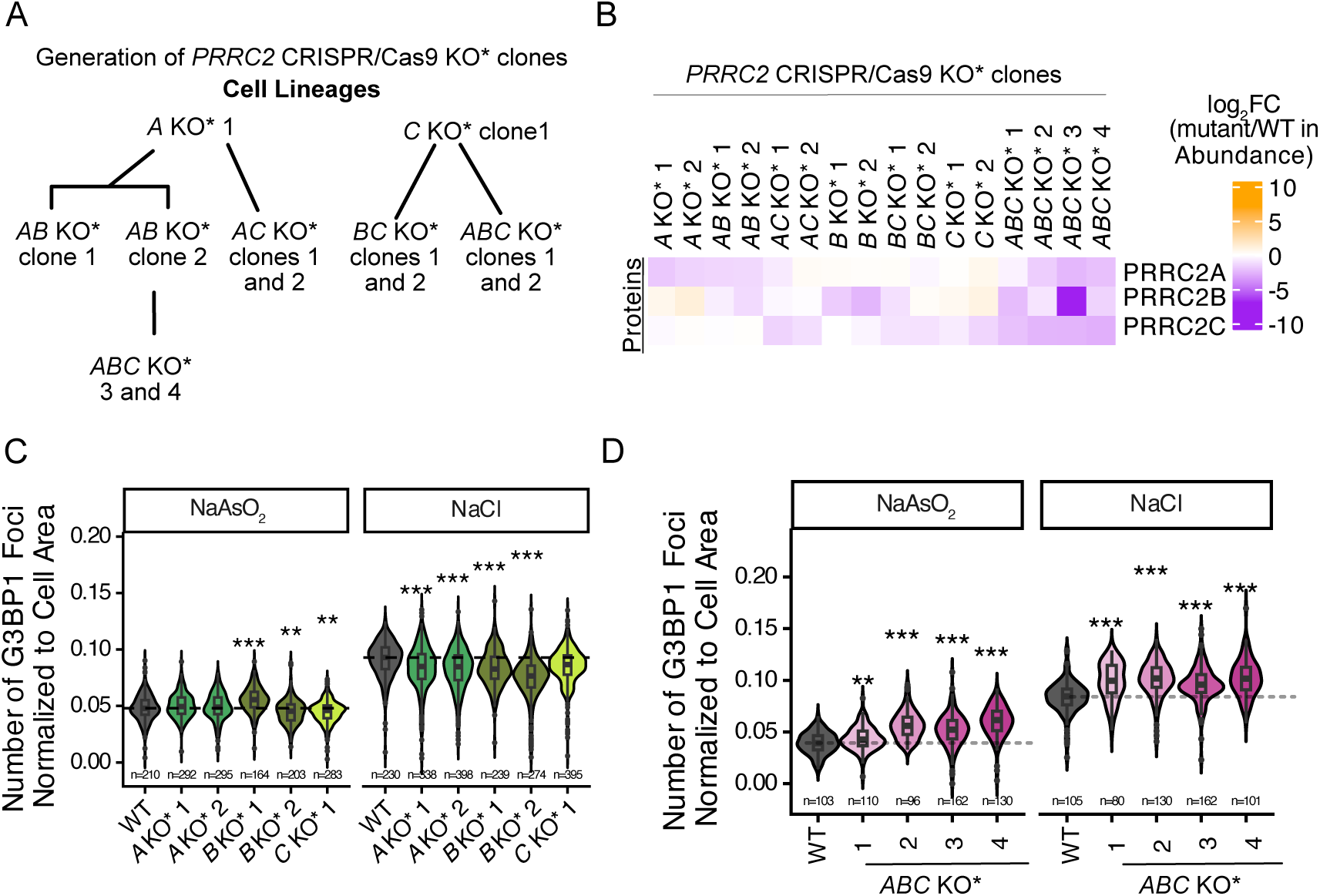
PRRC2 Proteins Promote SG Assembly in a Context-Specific Manner. **A)** Cell lineages for *PRRC2* CRISPR/Cas9 mutant generation. Single clones were isolated. The asterisk indicates that detectable, low-abundance peptides were found in the mass spectrometry analysis of the proteome. **B)** Heat map showing PRRC2 protein abundance log_2_ fold-change in all *PRRC2* mutants. **C)** Violin plots showing the number of G3BP1 foci normalized to cell area for single KO*s. Wilcox test with Holm correction, mutants compared to WT. * P-adjusted < 0.05, ** P< 0.01, *** P <0.001. The data show combined results from 3 biological replicates. n = total number of cells. The black dashed line indicates the median for WT. **D)** Violin plots showing the number of G3BP1 foci normalized to cell area for *ABC* KO* clones. Wilcox test with Holm correction, mutants compared to WT. * P-adjusted < 0.05, ** P< 0.01, *** P <0.001. The data show combined results from 3 biological replicates. n = total number of cells. The black dashed line indicates the median for WT.

Published work has shown that CRISPR Cas9-mediated editing can still produce functional N-terminal truncations or residual proteins due to weak nonsense-mediated decay responses, alternatively spliced variants or alternative translation start sites^85^. To validate total protein loss, we used mass spectrometry-based quantification of clonal cell lysates, which were subsequently used for total proteome analysis (**Fig. S6A,** methods) that showed high reproducibility across replicates (**Fig. S6B-C**). Quantification of PRRC2 proteins revealed that CRISPR Cas9-mediated editing led to the downregulation of PRRC2 protein abundances (**Fig. 6B, Table S7**). However, we still detected peptides upstream of the CRISPR-Cas9 cut sites in all clones and peptides downstream the cut site in one of the PRRC2A clones, which may have gone undetected in Western blots (**Fig. S6D-E**). Nonetheless, since the protein levels are downregulated, we chose to use these partial-knockout mutants and will refer to them as “KO*\**” clones.

Interestingly, this data revealed compensatory upregulation of other paralog(s) in single KO* cells, supporting their functional redundancy. Despite our attempts to use different sequential steps to obtain a triple knockout (**Fig. 6A**), no proliferating complete triple knockouts were obtained. However, we obtained four mutant clones with varying degrees of reduced PRRC2A, PRRCB, and PRRC2C protein levels (**Fig. 6B**), which we refer to as ‘*ABC* KO* clones’.

To determine whether PRRC2 proteins promote SG assembly, we quantified the number of G3BP1 foci in various single KO* mutants after acute exposure to oxidative or hyperosmotic stress using CellProfiler (35 min of 0.5 mM NaAsO_2_ or 200 mM NaCl, see Methods) (**Fig. S7A**). Most PRRC2 mutants showed reduced cell size (**Fig. S7B**), and changes in G3BP1 foci number reflected these cell size differences (**Fig. S7C**). Indeed, cell size was positively correlated with the number of SGs (**Fig. S7D**), necessitating cell-size normalization in SG quantification. Single PRRC2 KO* in general displayed a moderately positive role in SG assembly in stress specific fashion; SGs were reduced in *PRRC2B* KO* clone 2 and *PRRC2C* KO* clone 1 upon oxidative stress, while SGs were reduced in all tested single mutants under hyperosmotic stress (**Fig. 6C, Fig. S7A**). These defects were not due to reduced activation of the integrated stress response or to changes in G3BP1 protein abundance (**Fig. S7E**).

The subtle defects in SG formation observed in single mutants may be due to functional redundancy among PRRC2 proteins; therefore, *ABC* KO* clones were tested for defects in SG formation. We found that the number of SGs normalized to cell area unexpectedly increased in *ABC* KO* mutants under both oxidative and hyperosmotic stress. (**Fig. 6D, Fig. S8A**). This increase was not due to failed SG fusion, as the overall sizes of SGs were comparable between WT and mutants (data not shown). These defects were not attributed to changes in G3BP1 protein abundance or to increased ISR activation; rather, integrated stress response levels were diminished in *ABC* KO* mutants under hyperosmotic stress (**Fig. S8B**). Similar to the single mutants, the *ABC* KO* clones show reduced cell size, measured by cell area, which correlates with the extent of reduction in PRRC2 proteins’ abundances (**Fig. S8C, 6B**).

SGs are in dynamic flux with the P-bodies, because they share transcripts released from the translation pool ^6,86^. Given that mRNAs are the scaffolds of SGs and P-bodies, enhanced SG formation could imply that a larger fraction of transcripts shuttles to SGs than P-bodies in *ABC* KO* clones. To determine if *ABC* KO* clones showed altered P-body formation, the number of P-bodies was quantified using DCP1A as the marker during oxidative or hyperosmotic stress conditions. No significant changes were observed in P-body number, except for one or two *ABC* KO* clones (**Fig. S8D**), suggesting that increased SG formation cannot be explained by reduced partitioning of transcripts to P-bodies (note that the number of DCP1A foci did not correlate with cell area (**Fig. S8E**), so normalization to cell area was not applied).

We conclude that PRRC2 proteins have context-specific roles in SG assembly: PRRC2C promotes SG assembly under both test stress conditions, whereas PRRC2A and PRRC2B promote SG assembly only under hyperosmotic stress. In contrast to expectations, reducing the levels of all PRRC2 proteins increased SG formation, suggesting alterations in the mRNA pool or translation that seed SG formation.

### PRRC2 proteins redundantly promote translation required for cell growth

Functional redundancy amongst the PRRC2 proteins was tested using fitness as a phenotypic readout. Using PRRC2 mutants generated in HeLa Flp-In TREx cells, the growth rates of PRRC2 single, double and triple loss-of-function mutants were compared to WT using IncuCyte (see Methods). Given that increased PRRC2A, PRRC2B and PRRC2C transcripts were associated with various cancers^87^, growth rates of stable cells inducibly expressing each protein (3x-FLAG PRRC2A, OE-A; 3x-FLAG PRRC2B, OE-B; and 3x-FLAG PRRC2C, OE-C) were also assessed.

Relative to WT, all *ABC* KO* mutants showed a significant reduction in growth rate, suggesting that the PRRC2 paralog proteins redundantly promote cell growth (**Fig. 7A**). The strongest effect was observed in *ABC* KO* 4, exhibiting a 2-fold decrease in growth rate. The degree of growth defect correlated with the extent of reduction in PRRC2 proteins’ abundances. A significant reduction in growth rate was observed for one of the two *PRRC2A* KO* and *PRRC2C* KO* clones; clonal variability in the single mutants can partially be attributed to the compensatory upregulation of another paralog (**Fig. 6B**). No significant defects were observed for double mutants, suggesting the remaining PRRC2 paralog can compensate for function. Lastly, increasing the protein level of each PRRC2 protein did not affect the growth rate.

**Figure 7.**
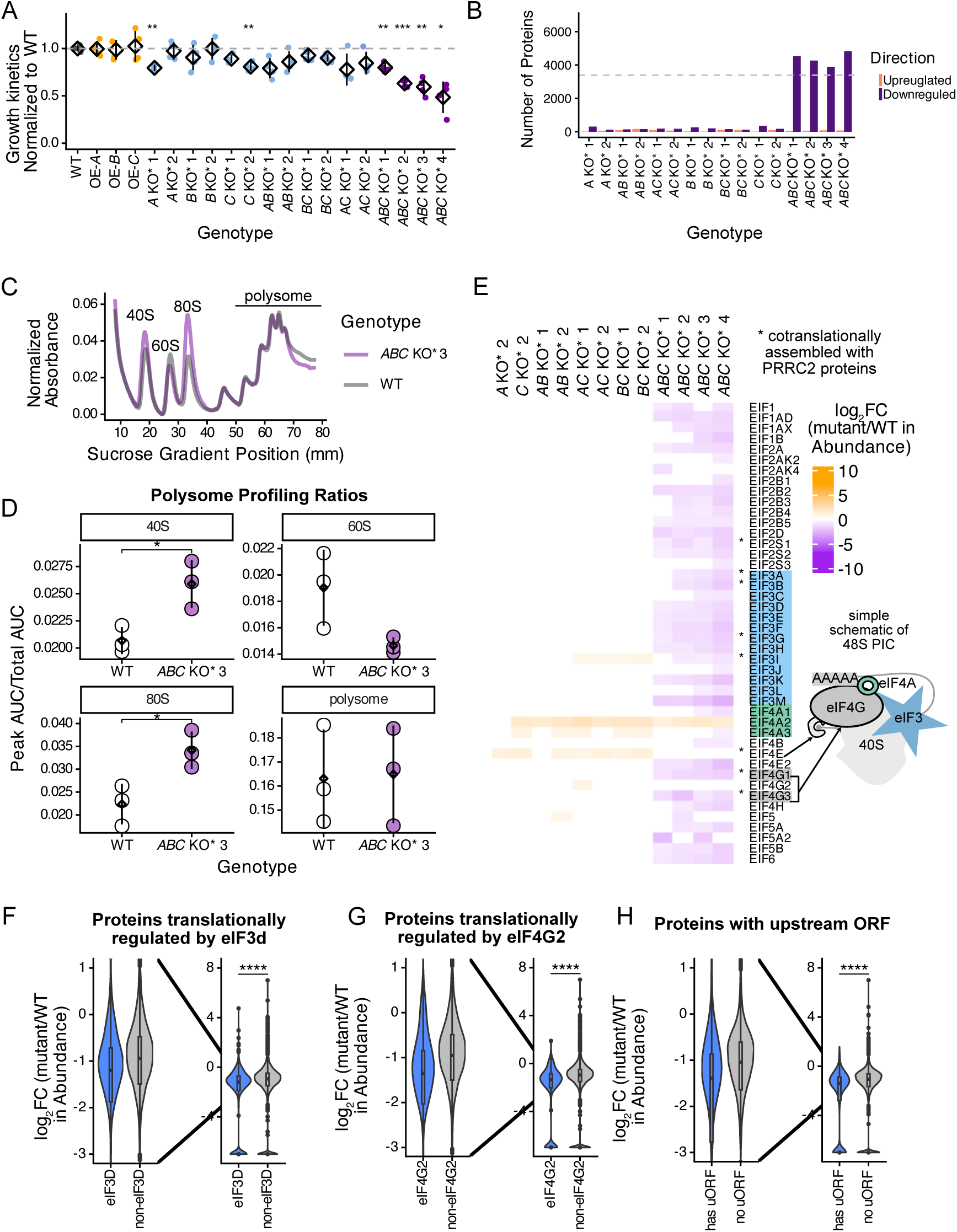
PRRC2 Proteins Function Redundantly in Promoting Cell Growth. **A)** Growth rate of *PRRC2* mutant cells. “OE” represents overexpression. Each dot represents the mean from three technical replicates. Black diamond and line represent the mean of four biological replicates and the standard deviation. The grey dashed line indicates the WT growth rate. Statistics for comparing mutant growth rates to WT were performed with Student’s t-test with Holm adjustment. *, P-adjusted < 0.05, **, P-adjusted < 0.01, *** P <0.001. **B)** Number of differentially expressed proteomes in *PRRC2* KO* clones. Grey dashed line = 50% of the detected proteome. **C)** Polysome profiling for WT and *ABC* KO* 3. The plot is a representative plot from one biological replicate. **D)** Plots showing the ratio of each peak normalized to the total AUC from polysome profiling experiments. Disomes were excluded from the polysome calculation. Black circle represents the mean from three biological replicates. The black line represents the standard deviation. Statistics were performed with Student’s t-test. * p-value < 0.05. **E)** Heatmap showing differentially expressed translation initiation factors. An asterisk indicates proteins are translationally assembled. **E)** Violin plot showing log_2_ fold-change for protein abundance in *ABC* KO* 3. Proteins were categorized as either eIF3d translation targets or non-eIF3d targets. Statistics were performed with the Wilcoxon rank-sum test. **** p-value < 0.0001. **F)** Violin plot showing log_2_ fold-change for protein abundance in *ABC* KO* 3. Proteins were categorized as either eIF4G2 translation targets or non-eIF4G2 targets. Statistics were performed with the Wilcoxon rank-sum test. **** p-value < 0.0001. **G)** Violin plot showing log_2_ fold-change for protein abundance in *ABC* KO* 3. Proteins were categorized based on the presence or absence of upstream open reading frames in their mRNA. Statistics were performed with the Wilcoxon rank-sum test. **** p-value < 0.0001

To understand the molecular functions of PRRC2 proteins, we analyzed the co-essentiality networks from the Cancer Dependency Map project (DepMap). DepMap contains CRISPR knockout screens across >1,000 cancer cell lines, providing information on common essential genes and cancer cell-specific genetic vulnerabilities^88^. Furthermore, the positive correlation in essentiality profiles across the cell lines (co-essentiality) identifies genes with the same pattern of dependency, revealing functional relationships between gene pairs (*i.e.,* gene-gene pairs’ co-essentiality in DepMap often reveals genes that encode functionally related proteins). Less frequently observed is a negative correlation between gene pairs, which often reveal negative regulatory relationships (e.g. MDM2 is negatively correlated with its substrate TP53 in the co-essentiality network).

Analyzing the top 100 co-dependent genes for each PRRC2 paralog from DepMap revealed the PRRC2 family of genes’ co-dependency to those involved in translation, ribosome biogenesis, cell death, or those encoding for mitochondrial proteins in either positive or negative direction (DepMap Public 25Q3+Score, Chronos, |correlations|>0.15) (**Fig. S9A, Table S8**). Notably, a co-dependent gene positively correlating with PRRC2A and PRRC2C is eIF4G2, a translation initiation factor involved in leaky scanning and translation and translation reinitiation^89^. This further implicates the role of PRRC2 proteins in translation. In addition, PRRC2C is negatively correlated with the tumour suppressor TP53, suggesting PRRC2C may have opposing functional roles in cell growth, consistent with PRRC2C’s positive role in cell growth (**Fig. 7A**).

Co-dependency analysis also indicates paralogs’ functional differences. PRRC2A is a common essential gene with 696/1186 tested cancer cell lines showing dependency. In contrast, only 58/1186 cancer cell lines were dependent on PRRC2C and none for PRRC2B. PRRC2A and PRRC2C are positively correlated with each other, while PRRC2B is not correlated with either paralog, suggesting that PRRC2A and PRRC2C are more functionally related among the three paralogs. Additionally, we find that KO of *PRRC2B* results in opposing phenotypes from the KO of *PRRC2A* or *PRRC2C* in a subset of genome-wide CRISPR KO screens performed in HAP1 cells^90^ (*i.e.,* positive genetic interactions between Gene A and PRRC2B, while negative genetic interactions are observed between Gene A and PRRC2A and/or PRRC2C; data not shown).

Given *PRRC2* genes*’* genetic co-essentiality and PRRC2 proteins’ interactions with translation factors, PRRC2 proteins’ function in translation was investigated by analyzing changes in proteome abundance and polysome profile in *ABC KO\** clones. Quantitative mass spectrometry analysis identified some significant changes in the single and double *PRRC2* mutants. In contrast, we observed proteome-wide reductions in the *ABC* KO* clones, with over 50% of the detected proteome expressed by 6775 genes showing significant decreases in abundance (**Fig. 7B**). This supports PRRC2 proteins’ redundant function in translation.

Polysome profiling of the *ABC* KO* clone with the most severe depletion, *ABC* KO* 3, displayed no changes in the polysomes fractions, indicating that translation elongation is not affected by the PRRC2 proteins (**Fig. 7C-D**). However, we observed significant increases in the 40S and 80S fractions (**Fig. 7C-D**), suggesting there are defects in translation initiation. The increase in the 40S may be attributed to defects in the joining of the 48S PIC with the 60S ribosome. The increase in 80S may be due to stalled monosomes or slower rates of translation initiation and is consistent with previous report on the effect of depleting PRRC2 proteins on translation^29^. Together, the polysome profiling revealed that PRRC2 proteins promote translation initiation.

A previous study reported that PRRC2 proteins are cotranslationally assembled with specific translation initiation factors, including eIF3b, eIF4G1 and eIF4E^29^. As a complex component, loss of PRRC2 proteins could destabilize the rest of the cotranslationally assembled complex and cause complex degradation, similar to what is known for complex stability ^91,92^. Indeed, the majority of the translation initiation proteins were downregulated to various degrees in the *ABC* KO*s (**Fig. 7E**). All cotranslational complexes, aside from eIF4E, were significantly downregulated, supporting the interactome data showing PRRC2 proteins are in a complex with components of the 48S preinitiation complex. Interestingly, we found one translation initiation factor, eIF4A2, was significantly upregulated in all four *ABC* KO* clones and the double mutants, suggesting a potential compensation mechanism. The decrease in translation initiation factors could be the primary cause for the global decrease in proteome abundance.

The co-essentiality analysis and protein interaction profiling suggest that PRRC2 proteins function together with eIF3 and eIF4G2 (**Fig. 3E, S9A**). If this is the case, the transcripts whose translation is regulated by eIF3 and eIF4G2 are expected to show decrease in protein abundance upon loss of PRRC2 proteins. We extracted the list of genes that were determined to be translationally regulated by eIF3d or eIF4G2 using polysome profiling^93,94^. All proteins quantified in the total proteome analysis were divided into two groups: those regulated by translation initiation factor and those that are not. Analysis of the protein abundance fold-changes between the two groups showed that the eIF3d and eIF4G2-regulated group shows significantly greater decrease in abundance in all four *ABC* KO* clones^93,94^ (**Fig. 7F-G, S9B-C**).

The PRRC2 proteins are implicated in the translation of transcripts containing upstream ORFs (uORF) and overlapping ORFs (oORFs), a function shared with eIF4G2 and eIF3 complex ^29,45–47,89,95^. Analysis of a ribosome profiling dataset^96^, which searched for genes with translation outside of the canonical coding region, revealed that transcripts containing uORFs were more significantly reduced in abundance in all four *ABC* KO* clones (**Fig. 7H, S9D**). The abundance changes suggest that loss of PRRC2 proteins preferentially affects the expression of uORF-containing transcripts. The effect of PRRC2 proteins on oORFs showed a more significant reduction in abundance in two out of four *ABC* KO* clones, while no significant differences were observed in other clones (**Fig. S9E**). This suggests that the loss of PRRC2 proteins primarily affects translation of uORF-containing transcripts.

In summary, our data support the idea that PRRC2 proteins redundantly promote cell growth through their shared functions in translation. The proteomics, DepMap co-dependency analysis and polysome profiling suggest PRRC2 proteins function in the translation initiation step with eIF3 and eIF4G2, with a bias for translating uORF-containing transcripts.

### PRRC2C interacts with eIF3 complex via its α helix

To determine how PRRC2 proteins promote translation by interacting with the eIF3 complex and eIF4G2 protein, we used deletion constructs lacking specific regions in PRRC2C and performed AP-MS analysis. Truncation constructs lacking the BAT2 domain, the putative coiled-coils, or the middle region containing the RG-motifs were expressed in HEK293 Flp-In T-REx cells, processed for AP-MS and analyzed in quantitative fashion (see Methods).

Overall, all three deletion mutants identified a larger number of high-confidence preys than the full-length PRRC2C protein (AvgP≥0.98) (**Fig. S10A**), despite comparable levels of expression (**Fig. S10B**). GO enrichment analysis of the AP-MS results showed that the mutants maintained their interactions with proteins involved in translation (**Fig. S10C, Table S9**), indicating that removing a single region is insufficient to completely abrogate PRRC2C’s interactions with translation factors. Notably, removing the BAT2 or the middle RG motif regions promoted novel interactions between PRRC2C and nucleoplasm proteins and those with transcription coregulator activity, suggesting that these regions regulate PRRC2C’s localization.

We hypothesized that removing the BAT2 region may enhance solvent exposure of PRRC2C’s SLiMs, which are important for nuclear localization or protein interactions. To test this hypothesis, we used the AlphaFlex workflow on PRRC2C full-length and PRRC2C-BAT2Δ and calculated the solvent accessible surface area of three SLiMs: two nuclear localization sequences (TRG_NLS_MonoExtN_4 and TRG_NLS_MonoCore_2) and an evolutionarily conserved SLiM for CtBP transcriptional co-repressor located C terminal to the BAT2 domain (LIG_CtBP_PxDLS_1_2; see Methods). PRRC2C-BAT2Δ displayed a higher solvent accessible surface area for LIG_CtBP_PxDLS_1_2, but only marginal differences for NLS SLiMs (**Fig. S10D-E**). The analysis indicates that removing the BAT2 region altered conformational dynamics and exposed an SLiM involved with nuclear protein interactions, with minimal effect on NLS. While this analysis supports PRRC2C-BAT2Δ’s increased interactions with nuclear proteins, this is unlikely to be due to increased accessibility of the NLS, which would promote PRRC2C-BAT2Δ’s nuclear localization.

Analysis of quantitative changes in the interactome revealed that many interactors had increased interactions with all deletion mutants, and a small subset of interactors decreased when the coiled-coils and middle RG motif region were removed (**Fig. S10F**). The decrease in interactors indicates that the interactions are mediated by these specific regions. For example, removing the middle RG motif region caused reduced interactions with five proteins, including TRA2A, TRA2B and SRSF9, which are involved in mRNA splicing (**Fig. S10G**).

The most notable change in interactions was a significant decrease in interactions with eIF3b, eIF3m, eIF3k and eIF4G2 upon removal of the putative coiled-coils (**Fig. 8A**). Other eIF3 subunits showed the same trend, while there was a more modest decrease in interactions with components of the eIF4F complex except for eIF4G3. These findings suggest that the putative coiled-coil region of PRRC2C facilitates interactions with the eIF3 complex and eIF4G2.

**Figure 8.**
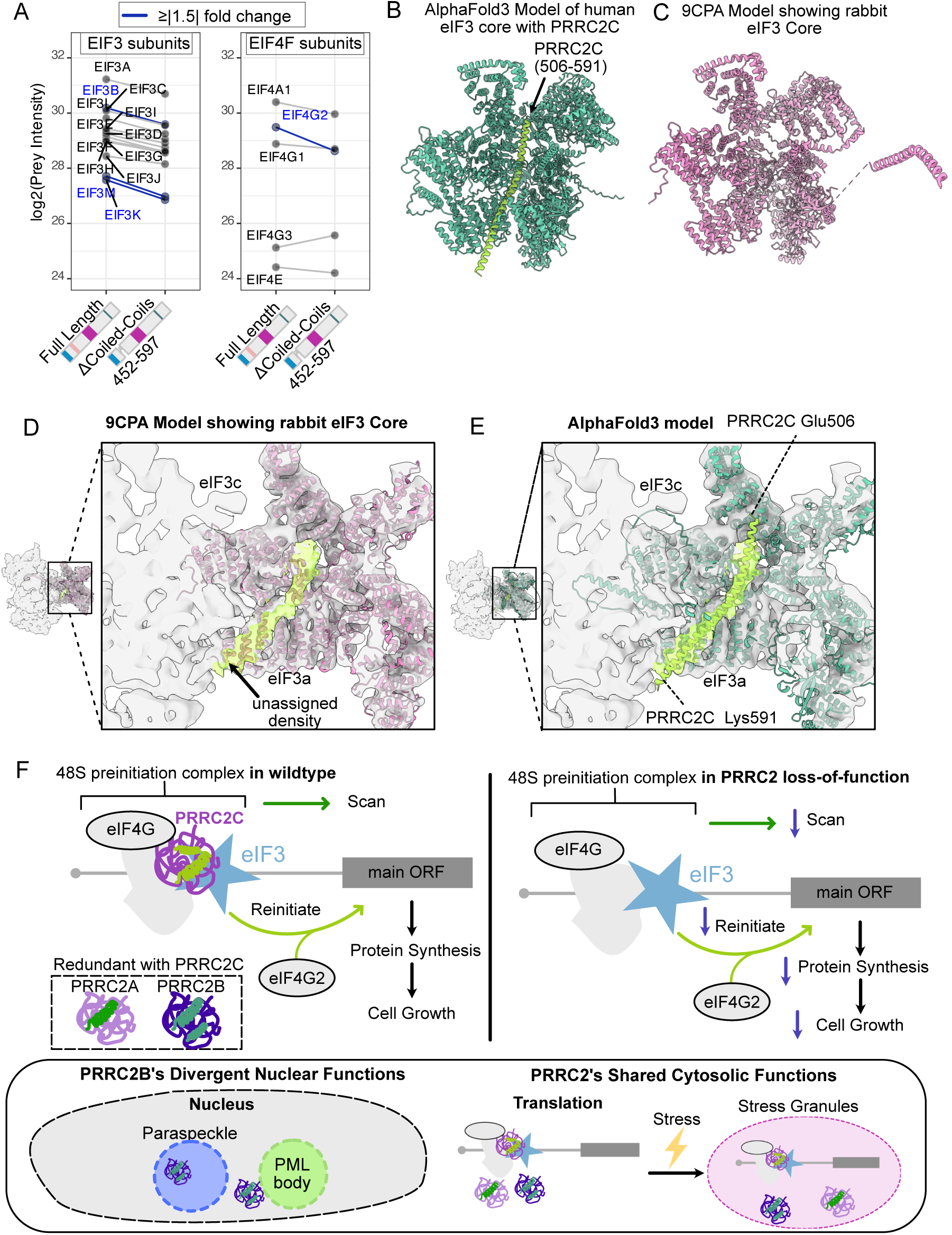
PRRC2C Interacts with eIF3 Complex through an α helix. **A)** Line plot showing log_2_ (Prey Intensity) in full length and PRRC2C-Δcoiled-coil. Only high confidence preys were displayed in the plot. ≥ 1.5-fold decreases are indicated in black with blue lines, and those below this threshold are indicated in gray. **B)** AlphaFold3 model of human eIF3 core with PRRC2C residues 506-591. The eIF3 core is coloured in green, while PRRC2C is coloured in bright green. The N-terminal 1-20 results for eIF3c and the C-terminus of eIF3a are not rendered for clarity. **C)** Experimental model of rabbit eIF3 core (PDB:9CPA).**D)** Cross-section of 43S preinitiation complex electron density with published atomic model. Unmodelled density is highlighted in yellow. Only the eIF3 core from the published model (PDB:9CPA) is shown. **E)** Cross-section of 43S preinitiation complex electron density with AlphaFold3 model. **F)** PRRC2 proteins redundantly promote translation with eIF3 to promote protein synthesis and cell growth. In PRRC2 loss-of-function mutants, the 48S preinitiation complex is less efficient in scanning and reinitiation. Consequently, this leads to decreased protein synthesis and cell growth. Upon stress, all PRRC2 proteins localize to Stress Granules. PRRC2B has divergent nuclear functions, including paraspeckles and PML body.

To investigate the interaction of PRRC2C’s putative coiled-coil domain with the eIF3 complex, we used AlphaFold3^97^ to model how the domain binds the eIF3 complex. The eIF3 core complex (a, c, e, f, h, k, l, and m) and PRRC2C coiled-coil region were used for modelling. The putative coiled-coil region in PRRC2C contains two α helices that were previously predicted to form a coiled-coil (**Fig. S9H**). Initial AlphaFold3 prediction suggested that the second α helix was responsible for PRRC2C’s interaction with eIF3a and eIF3c in the complex, so modelling was focused on those key residues (residues 506 to 591). The AlphaFold3 model predicts that this α helix from PRRC2C binds to a groove in the eIF3 core complex (**Fig. 8B**). This interaction has a high predicted template modelling score (pTM = 0.74), and a moderate interface predicted template modelling score (ipTM = 0.75), indicating the overall structure is predicted with high confidence while the positioning of the binding interface is moderately accurate. The AlphaFold3 model for the components of the core eIF3 complex closely resembled an experimental model determined from cryo-EM of the mammalian 43S preinitiation complex^98,99^ (PDB ID: 9CPA) (**Fig. 8C**). Inspection of the cryo-EM density map (EMDR: emd3057) and the 9CPA model revealed an unassigned α-helical density in the eIF3 core adjacent to the eIF3a and eIF3c subunits (**Fig. 8D**). Remarkably, aligning the AlphaFold3 model with the published model showed the unassigned density was filled by the PRRC2C α helix (**Fig. 8E**).

In summary, protein-protein interaction domain mapping identified the key region in PRRC2C that facilitates interactions with the eIF3 complex. AlphaFold3 modelling predicted that an α helix within the putative coiled-coil region of PRRC2C interacts with eIF3a and eIF3c. The α helix fits into an unassigned α-helical density observed in previously published cryo-EM density of the mammalian eIF3 complex, providing a strong independent support for the accuracy of the AlphaFold3 model.

## DISCUSSION

Elucidating the functions of intrinsically disordered proteins remains challenging due to their lack of stable tertiary structure and the conformational heterogeneity that underlies their dynamic molecular interactions. The PRRC2 proteins represent an evolutionarily conserved family of largely intrinsically disordered paralogs whose molecular functions remain poorly defined. Our integrative analysis, combining proteomics, cell biology, genetics, and computational modeling, identified PRRC2 proteins as translation regulators that associate with the 48S PIC to promote translation in basal conditions.

Despite their overall disordered nature, PRRC2 paralogs share conserved architectural features. While sharing many compositional features and evolutionarily conserved molecular features, we also observed variations amongst the paralogs, suggesting both conserved molecular functions alongside paralog-specific adaptations. Notably, PRRC2C displays compositional divergences within the BAT2 region (**Fig. 1D**). This is consistent with our structural and interaction data showing PRRC2C’s interactions with the translation machinery via the coiled-coil region, rather than the BAT2 region reported for PRRC2A and PRRC2B ^31,32^. These findings highlight how intrinsically disordered proteins can maintain conserved functional outputs (*i.e.,* interaction with translation factors), while evolving distinct molecular interfaces. The bioinformatic analyses provided additional descriptive features of PRRC2 proteins, which can be interrogated for their contribution to PRRC2 function in the future.

Protein-protein interaction mapping revealed that the PRRC2 proteins are closely integrated with the translational apparatus at the basal state, showing strong enrichment for translation initiation factors, ribosomal proteins, and RNA regulatory complexes (**Fig. 3**). These interactors suggest that PRRC2 proteins are not merely stress-responsive factors, but active components of basal translation regulation. Under stress conditions, their interaction networks undergo extensive remodeling, involving SG components, translation initiation factors, and ribosome quality control factors. This dynamic rewiring supports a model in which PRRC2 proteins function at the interface of translation initiation and stress-responsive translational reprogramming, which warrants future studies.

Mechanistically, our structural and protein-protein interaction analyses indicate that PRRC2C directly interacts with the eIF3 complex, specifically eIF3a and eIF3c, via an α helix within its predicted coiled-coil region (**Fig. 9E**). Given the highly similar proteomics interaction profiles and redundant phenotypes observed across paralogs, we propose that all three PRRC2 proteins associate with the 48S PIC through interactions with the translation initiation factors (*e.g.* eIF3 complex and eIF4G), thereby facilitating translation initiation. This model is further supported by proteomic signatures showing preferential dysregulation of proteins encoded by transcripts regulated by eIF3d and eIF4G2, as well as transcripts containing uORFs upon loss of three paralogs. Together, these observations suggest that PRRC2 proteins contribute to translation initiation.

Genetics revealed the apparent functional redundancy among paralogs. While depletion of individual PRRC2 proteins produced modest phenotypes, combined depletion resulted in severe defects in cell growth, reduced global proteome abundance, and altered SG formation. The reciprocal upregulation of remaining paralogs upon loss of one family member suggests compensatory buffering, potentially through transcriptional adaptation, translation control, or altered protein stability. Such buffering may obscure paralog-specific functions under normal conditions while preserving essential translational capacity.

Our identification of PRRC2 proteins as translation-associated factors supports a new framework for understanding SG function. Rather than remaining in a translationally dormant state, SGs and SG proteins increasingly appear to be involved in translational reprogramming during stress^16,100^. The association of PRRC2 proteins with eIF3 complex also implicates their shared function in maintaining selective translation during stress, in line with a recent study showing that SGs support ISR-driven translational reprogramming^16^. Corroborating with this idea, others have shown that PRRC2 proteins are important for ATF4 translation in ER stress, the primary downstream effector of the ISR^29^. Furthermore, emerging data implicating PRRC2 proteins in the recognition of m^6^A-modified transcripts raise the possibility that they may selectively promote translation of methylated stress-responsive mRNAs, analogous to m^6^A-dependent recruitment of eIF3 to specific Hsp70 transcripts^27,35,36,38,73,101,102^.

The translational functions of PRRC2 proteins may also have implications for cancer biology. PRRC2 proteins are elevated across multiple cancer types^103^, and cancer-associated mutations in eIF4G2 disrupt its interactions with PRRC2 and impair IRES-dependent translation initiation^104^. These observations suggest that PRRC2 proteins may contribute to oncogenic translation control, although direct functional evidence will be required to establish their role in tumour progression.

In summary, our work identifies PRRC2 proteins as components of the translation initiation machinery that redundantly promote translation and cell growth while linking translational control to stress responses (**Fig. 9F**). Among the paralogs, PRRC2B additionally exhibits expanded interactions with nuclear RNA bodies, suggesting functional diversification. Structural evidence of direct interaction between PRRC2C and the eIF3 complex provides a mechanistic framework for understanding the regulation of translation by SG proteins. Beyond defining the roles of PRRC2 proteins in translation control, stress granules, and potentially in translation reprogramming, this work highlights a generalizable strategy for functionally dissecting intrinsically disordered proteins.

### LIMITATIONS OF THIS STUDY

We acknowledge that the low overlap in interactors identified between AP-MS and BioID data may partly be due to potential post-lysis interactions identified in AP-MS data.

## METHODS

### Cell Culture

Cell lines were regularly monitored for mycoplasma contamination. HEK293 Flp-In T-REx (Invitrogen, R780-07) and HeLa Flp-In T-Rex (Arshad Desai Lab) cell lines were cultured in Dulbecco’s modified Eagle’s medium (DMEM) high glucose (Wisent, 319-005-CL) supplemented with 5% fetal bovine serum, 5% calf serum and penicillin/streptomycin. HAP1 cells were cultured in Iscove’s modified Dulbecco’s medium (IMDM) (Wisent, 319-210-CL) supplemented with 10% fetal bovine serum and penicillin/streptomycin.

### Flp-In Stable Cell Line Generation

Expression clones were generated with Gateway cloning (Thermo Fisher). Entry clones with validated RefSeq sequences (PRRC2A NP_542417.2 (UniProt P48634), PRRC2B NP_037450.2 (UniProt Q5JSZ5), PRRC2C NP_055987.2 (UniProt Q9Y520-4) were synthesized by Gene Universal and were recombined into destination vectors miniTurbo-3×FLAG or 3×FLAG (gifts from Anne-Claude Gingras, Lunenfeld-Tanenbaum Research Institute). Destination clones and pOG44 (Flp-In Recombinase) were co-transfected into Flp-In TREx cells using JetPrime transfection reagent (Polyplus, CA89129-924). After 24 hours of transfection, cells were passaged and allowed to recover for 24 hours. Then, selection was applied using growth media supplemented with Hygromycin B (200 µg/mL; Wisent, 450-141-XL) until clonal colonies were visible. Colonies were pooled for experiments.

The expression of miniTurbo-PRRC2C was 3∼4 times lower than its paralogs, PRRC2A and PRRC2B (based on the recovery of self-biotinylated PRRC protein level), which likely contributed to poor BioID labelling and a low number of high-confidence preys. We attempted to express miniTurbo-PRRC2C in *PRRC2C* KO* cells to ameliorate expression, but there was no significant improvement.

### Generation of endogenously fluorescently tagged HAP1 cells using CRISPR-Cas9

HAP1 cells (Horizon Discovery) were endogenously tagged by following the protocol from Xiao *et al.* ^105^. Homology arms were synthesized by Twist Bioscience and cloned into the donor plasmid (gift from Jason Moffat). Plasmids containing the guide RNA (pX459) and the donor template were electroporated into cells, using Amaxa Nucleofector (program Y-005) and Nucleofector kit R (VCA-1001). After 24 hours of transfection, cells were passaged and allowed to recover for 24 hours. Then, selection was applied using growth media supplemented with puromycin (2 µg/mL; Biobasic, 400-160-EM) until clonal colonies were visible. Colonies were pooled for experiments.

### PRRC2 Knockout Generation and Genotyping

PRRC2 mutant cell lines were generated using the CRISPR-Cas9 system. Single guide RNAs (sgRNA) targeting N-terminal of *PRRC2A*, *PRRC2B* and *PRRC2C* was cloned into a pX459 plasmid. Then, the pX459 plasmid was transiently transfected into HeLa Flp-In TREx cells with JetPrime transfection reagent. Puromycin selection was applied for 24 hours after transfection. Then, cells were allowed to recover for at least 24 hours. Single clones were isolated by serial dilution and seeding into a 96-well plate. After each clone was expanded, the genomic DNA was extracted using GeneJET Genomic DNA Purification Kit (K0721). PCR was performed around 500 base pairs upstream and downstream of the cut site, and the product was sent for Sanger sequencing at The Centre for Applied Genomics, The Hospital for Sick Children. Indels were confirmed using Inference of CRISPR Edits^106^ (ice.editco.bio).

### IDR analysis of PRRC2 proteins

Disorder scores were generated using Metapredict V3.0^107,108^ (metapredict.net). A default threshold of > 0.5 was considered intrinsically disordered. For the prediction of folded structure, the PRRC2 sequences were submitted to the AlphaFold3 server^97^, and the pLDDT scores from the best-ranked were used for the plot. The coiled-coil probabilities were calculated using Marcoil^109,110^. The probability of a prion-like domain was predicted using PLAAC with a default core size of 60 and using Homo sapiens frequency^111^. The compositional Z-scores from NARDINI+ were calculated using the “NARDINI+_From_FASTA” Google Colab notebook (https://github.com/kierstenruff) ^57^.

For the analysis of human PRRC2 proteins, RG motifs were identified using regular expression, as described in Thandapani *et al.*^60^. The nuclear localization signals were found using the ELM SLiM database (elm.eu.org)^112^. The PY-NLS was identified by regular expression in ScanProsite (prosite.expasy.org)^113^. As described in Lee *et al.*, the NLS was counted as 50 amino acids (beginning 40 residues N-terminus of the PY to 10 residues C-terminus of that motif) using regular expression^114^.

The canonical sequences from PRRC2 orthologs were extracted from Ensembl^115^. Then, the sequences were aligned using MAFFT (v7.511) with default settings^116^. The phylogenetic tree was constructed in the MAAFT web server using neighbour joining and default methods (model = JTT) with 1000 bootstrap. The sequences were searched in InterPro to determine whether the BAT2 domain was present. A predicted coiled-coil was considered present if it was predicted with high confidence (probability >90). The phylogenetic tree was plotted in R using the ggtree package. Heatmaps were aligned to the phylogenetic tree using the aplot package.

### Conformation Ensemble Analysis using AlphaFlex

Using the AlphaFlex^53^ definition of an IDR, PRRC2A, PRRC2B, PRRC2C, and PRRC2C-ΔBAT2 were all predicted to be fully disordered. IDPConformerGenerator (V0.7.19) was used to generate 10,000 conformations for each protein sequence for analysis^117,118^. The secondary-structure propensity was derived from sequence statistics as observed in the annotated IDPConformerGenerator database from non-redundant RCSB PDB^119^ X-Ray crystal structures of at least 2.0 Å. Global structural properties (radius of gyration, hydrodynamic radius, end-to-end distance, asphericity, and global surface accessible surface area) were analyzed using the same methods described in the AlphaFlex study^53^.

Short-linear motifs were identified for each sequence using the Eukaryotic Linear Motif (ELM) webserver^112^. Solvent accessible surface area for patches of residues was calculated using the FreeSASA^120^Python library (V2.2.1) from PyPi. Residue patches that were categorized as surface accessible needed to have a combined surface area that is at least 20% of the maximum allowed solvent accessibilities of each residue type^121^.

### Evolutionary Signature Analysis

The evolutionary signatures were computed in the same way as Pritišanac et al. ^63^(https://github.com/IPritisanac/IDR_ES). However, we restricted our analysis to 1-to-1 ENSEMBL orthologs, because inaccurate assignment of orthologs can confound the results. Additionally, absolute values for Z-scores were capped at 20 for easier plotting and visualization.

### DepMap Co-Dependency Network

The top 100 co-depedent genes were extracted from the DepMap portal for PRRC2 proteins (https://depmap.org/portal)^88^. The DepMap Public 25Q3+Score, Chronos gene correlations were filtered for absolute correlations ≥ 0.15 and then visualized in Cytoscape(V3.10)^122^.

### Lattice lightsheet Live Cell Imaging

HAP1 cells were seeded into 4-well Nunc™ Lab-Tek™ II Chambered Coverglass (Borosillicate Glass 1.5) 18 hours prior to imaging. About two hours prior to imaging, cells were washed once with D-PBS. Media was replaced with Iscove’s Medium, without phenol red (Wisent, 319-210-CL) supplemented with 10% FBS and penicillin/streptomycin.

Live cell imaging was performed on a ZEISS Lattice Lightsheet 7 microscope (44.83x/1.0 water immersion). Cells were kept at 37°C with 5% CO2 and 70% humidity. The stress solution was applied 5 min before acquisition. Then, a time lapse was acquired for a total of 10 min with 30 s intervals. X-stacks were acquired deskewed in Zen software (Zeiss).

### Immunofluorescence microscopy

Cells were plated on sterile coverslips in 12-well plates and allowed to adhere overnight. After appropriate experimental treatment, cells were washed once with PBS and fixed with 4% paraformaldehyde in PBS for 10 min at room temperature. Post fixation, cells were washed twice with TBST (0.1% Tween-20). Cells were permeabilized with 0.1% TritonX-100 in TBST for 10 min, washed twice with TBST, then blocked at room temperature with 5% skim milk or 5% BSA. Coverslips were moved to a humidified chamber for overnight staining at 4°C. The next morning, coverslips were washed thrice with TBST before incubation with Alexa fluorophore-conjugated secondary antibodies for 1 hour at room temperature. Then, cells were washed twice before incubation with DAPI 1µg/mL for 5-10 min at room temperature, followed by two washes with TBST, then with miliQ water before mounting with Prolong Gold. Coverslips were allowed to cure for at least 24 hours before imaging.

Immunofluorescence confocal microscopy was performed on either a Leica SP8 (HCPL APO CS2 63x/1.30 glycerol immersion, HCPL APO CS2 63x/1.40 oil immersion or HCPL APO CS2 40x/1.10 water immersion) or a Leica Stellaris 5 (HCPL APO CS2 63x/1.40 oil immersion). Z-stacks were acquired with either line, frame or stack sequential with bidirectional acquisition at a scan speed of 400 Hz. Pixel sizes were at a minimum of 1024×1024 pixels. Leica SP8 is equipped with PMT and HyD detectors while the Stellaris 5 used HyD S detectors.

Immunofluorescence images in lightning mode were acquired on a Leica SP8 microscope (HC PL APO CS2 100x/1.40 oil immersion). Images were acquired with stack sequential, bidirectional scanning and with Lightning adaptive deconvolution (refractive index set to 1.47). The pinhole size was changed to 0.2 AU. Samples were illuminated with a white light laser (499 nm and 553 nm) and detected with HyD detectors. The LAS X software (Leica, 3.5) was used to perform acquisition.

### Confocal Image Analysis

Line-scan analysis and movies were performed in FIJI(2.16). Line-scan analysis was performed by drawing a line using the arrow tool. The measured intensity was normalized to the mean grey value for plotting.

Three separate pipelines were created for analyzing images to adjust for differences in pixel sizes. All Z-stacks were compressed into maximum projects for analysis in CellProfiler (v4.2.8)^123^. Details are below.

For analysis of lightning confocal images, the module MeasureColocalization was used to measure the Pearson correlation coefficient and Manders’ coefficient across the entire immunofluorescence image. Pearson correlation measures whether there is a relationship between pixel intensities between two channels. Mander’s coefficient measures the cooccurrence by calculating the fraction of pixels from the first channel that overlap with pixels from the second channel. To quantify the size and shape of SGs, SGs were first segmented using IdentifyPrimaryObjects module using a global, robust background method. SGs with a diameter < 0.5 mm were excluded to minimize false positive segmentation. The diameter and form factor of each SG were measured using MeasureObjectSizeShape module. Pixels were converted into mm using the known pixel per mm size in image metadata.

For counting G3BP1 and DCP1A foci per cell, DAPI staining was used to identify cells. The cytoplasm was linked to each nucleus and segmented using IdnetifySecondaryObjects using propagation, global thresholding (minimum cross-entropy). MeasureObjectSizeShape was applied to the segmented cytoplasm (marked by G3BP1 signal) to determine cell area. Dead cells were filtered out based on the size of the cytoplasm. To enhance segmentation, the EnhanceOrSuppressFeatures module was applied to enhance speckle features. Then, SG segmentation was performed on the enhanced image using the IdentifyPrimaryObjects with an adaptive thresholding strategy (robust background). PBs were similarly segmented by using an enhanced imaged and a global thresholding strategy (robust background). A size minimum of >0.5 µm was included for PBs and SG segmentation. SGs and PBs were linked back to each cell using the RelateObjects module. Pixels were converted into mm using known pixel per mm size in image metadata.

### Sample Collection and Processing for BioID

PRRC2-miniTurbo-3×FLAG bait and negative controls (miniTurbo-3×FLAG-eGFP and miniTurbo-3×FLAG alone, parental cell line) in HEK293 Flp-In T-REx were seeded in a 10 cm plate. Bait expression was induced for 24 hours using 1µg/mL tetracycline in biotin-depleted media at 70-80% confluency. Biotin labelling was performed for 30 min using growth media supplemented with 50 µM biotin in three different conditions as described in **Fig. 4A**. Cells were washed with D-PBS and lifted using 0.05% trypsin. Cells were spun down at 500g for 4 min and resuspended in D-PBS. Cells were transferred to a pre-weighed Eppendorf tube, pelleted and frozen with dry ice. Sample processing and peptide collection are described in Tsai et al. Cell lysates were normalized within each bait group. Peptides were resuspended in 5% formic acid, and 1/12 of the total material was analyzed in Exploris 480.

### Sample Collection and processing for Affinity purification coupled to mass spectrometry

PRRC2-3×FLAG bait and negative controls (3×FLAG-eGFP and parental cell line) in HEK293 Flp-In T-REx expression were seeded into a 15 cm plate. Bait expression was induced for 24 hours using 1µg/mL tetracycline at 70-80% confluency. Cell pellet collection was identical to BioID samples. The frozen pellet was resuspended in ice-cold TAP lysis buffer (50 mM HEPES-NaOH pH8, 100 mM KCl, 2 mM EGTA, 0.1% NP-40, 10% glycerol, 1.5 mM MgCl2, with freshly supplemented inhibitors plus 1 mM PMSF and 1 mM DTT) at 4:1 (volume: weight). Cells were lysed by freeze-thaw and sonicated on ice for 30s(10s on, 5s off for three cycles) at 25% amplitude at 4°C. Then, 250 U (1 µL) of Benzonase was added to each tube, which was rotated (end-over-end) at 4°C for 20 min. Lysate volumes were normalized across all baits and were centrifuged at 20,817x g for 20 min at 4°C, and the supernatant was transferred to a fresh tube. Then, 12.5µL of 50% FLAG-M2 beads (Sigma Aldrich, M8823-5ML) was incubated with the lysate for 3 hours at 4°C using gentle end-over-end rotation. Washes were performed using a magnetic rack. Briefly, 1mL modTAP lysis buffer was used to resuspend the beads (up and down 4x) and transfer to a fresh 1.5mL tube, followed by one wash with FLAG rinse buffer (20mM Tris-HCl and 2 mM CaCl2) by resuspending the beads (up and down 4x). On-bead digest was done in 5 µL of 1µg of trypsin (in 20mM Tris, pH 8) overnight at 37°C. An additional 0.5µg of trypsin in 2.5µL of 20 mM Tris-HCl was added and incubated at 37 °C for 4 hours. Collected peptides were acidified to a final concentration of 2% formic acid, and 1/10 of the total material was used for DDA and DIA analysis.

### S-Trap sample collection and processing for total proteome MS analysis

Wildtype HeLa Flp-In T-REx and *PRRC2* KO* cells were collected from 12-well plates by trypsinization. Cell pellets were washed once with D-PBS before flash freezing on dry ice. Pellets were lysed in lysis buffer (5% SDS, 50mM TEAB pH 8.5, 2mM MgCl_2_) with a ratio of 15 mg pellet per 23 µL lysis buffer volume. Then, 100 units (0.4 uL) of Benzonase was added to each tube and incubated at room temperature for 5 min. Lysates (23 µL) were added to a new tube and spun down at 13,000g for 8min. The cleared lysate was transferred to a fresh tube. Then, 1 µL of reductant (5mM TCEP) was added to the cleared lysate and incubated at 55°C for 15 min. Then, 1 µL of alkylator (final 20 mM MMTS) was added to the cleared lysate and incubated at room temperature for 10 min. Next, 2.5 µL of acidifier was added (27.5% phosphoric acid, final ∼2.5% phosphoric acid). 165 µL of binding/wash buffer (100mM TEAB in 90% Methanol) was added to the cleared lysate and applied to a S-Trap micro spin column (ProtiFi). The S-trap was spun down at 4,000 g for 30s to trap proteins. Washes: 150 µL of binding/wash buffer, centrifuged at 4,000 g for 30s – repeated 3 times. The S-trap was centrifuged for 1 additional min to remove residual buffer. S-trap was transferred to a clean 1.5 mL tube for trypsinization. Trypsinization was performed in a water bath heated to 47°C for 1.5-2 hours. Proteins were digested with 10 µg of trypsin in a solution of 50mM TEAB (total 20uL digestion buffer). Then, 40 µL of elution buffer 1(50mM TEAB in water), buffer 2 (0.2% formic acid in water) and buffer 3 (50% acetonitrile in water) were applied sequentially and spun down at 4,000g for 1 min. Peptides were dried in a Speedvac and resuspended with 160 μL of 5% formic acid. Peptide concentration was measured using Pierce™ Quantitative Peptide Assays & Standards (Thermo Fisher, 23275). A total of 400 ng of peptides was analyzed in the Exploris 480.

### Liquid Chromatography and Mass Spectrometry

Peptides were separated in an Easy-nLC 1200 liquid chromatography system (Thermo Fisher). Samples were injected into a 10.5 cm 75 µm ID emitter tip, packed in-house with 3 µm ReproSil Gold 120 C18 (Dr. Maisch HPLC GmbH). Peptides were eluted at 200 nL/min using a 120 min gradient from 2.4% acetonitrile to 35% acetonitrle in 0.1% formic acid. The acetonitrile concentration was increased to 80% over an 8 min gradient and held for 16 min at 200 nL/min, for a total run time of 144 min. Samples were analyzed with an Orbitrap Exploris 480 (Thermo Fisher Scientific). The instrument was operated in positive ion mode with a sprat voltage of 2.4 kV, and an ion transfer tube temperature of 300 °C. Instrument parameters were set up using Thermo Xcalibur software.

Data-dependent acquisition was used to acquire data for generating a spectral library. MS1 scans (390–1010 m/z) were acquired at 120,000 resolution with a 3s cycle time and a normalized AGC target of 125%. Precursors with charges 2+ to 7+ and intensity >1×10³ were selected for fragmentation (NCE 33%) with 9 s dynamic exclusion. MS2 scans were acquired at 15,000 resolution, normalized AGC 400%, and maximum injection time 50 ms.

Data-independent acquisition was used to collect data for BioID, AP-MS and whole proteome analysis. Two alternating precursor isolation schemes (400–1,000 and 396–1,004 m/z) were applied. MS1 scans were acquired in centroid mode at 60,000 resolution (390–1,010 m/z, normalized AGC 100%, 55 ms maximum injection time). MS2 spectra were collected using 75 and 76 DIA windows of 8 m/z each at 15,000 resolution (145–1,450 m/z, normalized AGC 1000%, 23 ms maximum injection time; normalized collision energy 33%).

To generate a library for whole proteome analysis, tryptic peptides from wildtype HeLa Flp-In T-REx was collected using S-trap micro spin columns. Then, gas phase fractionation DIA was performed using six serial injections (400 ng peptides for each injection). A narrow, staggered precursor isolation window was used (4 m/z at 60,000 resolution, AGC target of 1000%, maximum injection time 55 ms, 33% normalized collision). The six injections covered 398-502, 498-602, 598-702, 798-902 and 898-1002 m/z range.

### Spectral Library Generation

Spectral library generation was performed as described in Tsai et al^70^. MS data included for spectral library generation were fractionated HEK293 lysate DDA data and PRRC2 AP-MS and PRRC2 BioID DDA data (HEK293+PRRC2 library).

The gas phase fractionation library was generated as described in Tsai et al. The only change was fixed modifications for Cysteine. We changed the fix modification to MMTS modification (+45.98772). Subsequent library optimization is identical to HEK293+PRRC2 library.

### DIA-NN Analysis of DIA MS data

The MS Thermo RAW files acquired in DIA and gas phase fractionation was demultipexed in MSConvert (ProteomeWizard;V3) with “peakpicking vendor msLevel-1” and “demultiplex optimization = overlap_only mass error=10.0ppm”. The demultiplexed MzML files for BioID and AP-MS were analyzed in DIA-NN (V1.8.1) with the HEK293+PRRC2 library. For analysis of BioID runs, peptides with alkylated cysteines were filtered out of the library to reduce false positives. DIA-NN was configurated with match-between-runs enabled, normalization was turned off and “--report-lib-info” was flagged to generate a fragment report (report.tsv).

### BioID and AP-MS MS data processing and SAINTq

The fragment level (MS2) data was extracted from the fragment report. Fragments that passed Lib.Q.Value ≤ 0.01 and Lib.PG.Q.Value ≤ 0.01 were used for further analysis. Median intensity normalization was applied across all samples. SAINTq analysis was performed as described in Tsai *et al.* using the normalized intensities. SAINTq files were visualized as dotplots in ProHits-viz^124^ (https://prohits-viz.org/).

### Total Proteome Data Analysis

Demultiplexed DIA files were analyzed in DIA-NN using the HeLa Flp-In gas phase fractionation library. The DIA-NN settings were the same as above, aside from the addition of one command “–fixed-mod UniMod:39,45.987721,C” for peptides containing MMTS-modified cysteines.

Statistical analysis on the proteome was performed using MSStats (V4.16). The fragment report file (report.tsv) from DIA-NN was converted and filtered for MSStats using MSstatsConvert. All settings for MSstatsConvert were kept at default aside from one setting – proteins with 1 feature were removed. Fragments were summarized into protein level using dataProcess function with default settings. Briefly, feature intensities were normalized using medians across samples. The default summarization method, Tukey’s Median Polish, was performed using all features and imputation was enabled. Pairwise comparisons for differential protein abundance analysis were performed using groupComparison function. Proteins with p-adjusted level ≤ 0.05 and log2 fold-change ≥ 0.58 (1.5 fold change) were considered significant. Genes with –Inf/Inf were replaced with –7/7 for plotting.

For PRRC2 peptide log2 fold-change calculations, the log2 feature intensities matching the same stripped peptide sequence was summed for each replicate. Then, the mean intensity from three replicates was used for fold-change calculations. Then, the fold-change was determined by log2(mutant intensity)-log2(WT intensity).

### Cell growth Assay

Cells were seeded in a 96 well plate (Corning 3603) at a density of 1.5 × 10^3^ cells per well. All cell lines were plated in duplicate wells as technical replicates within the same plate. After cells had settled for 4-6 hours, the media was changed to media containing 1 µg/ml tetracycline for wells containing overexpression cell lines. Overexpression cell lines were generated by introducing the 3×FLAG-tagged PRRC2 destination clones into HeLa Flp-In T-REx so that cells will express both endogenous and 3×FLAG-tagged constructs. Cells were given fresh media 1 hour before imaging. For the overexpression cell lines, the growth media were supplemented with fresh tetracycline 1 hour before imaging.

After 24 hours of seeding, the plate was placed into a Sartorius Incucyte (V2023). Each well was imaged every 4 hours over a period of 3 days under phase contrast at a 20x objective. The confluency measurements were extracted using a basic analyzer in Incucyte software. Then, using the growthrates package in R, a spline was fitted to each growth curve, and the growth rate (mumax) was extracted. The mean growth rate from the technical replicates was plotted. One outlier technical replicate from a biological replicate was removed. For easy comparison to wildtype, the growth rate was normalized to the mean wildtype growth rate.

### Polysome Profiling

Cells were cultured in 15-cm dishes until 95% confluency. The growth media (DMEM) was refreshed for 30 min prior to collection. Cycloheximide (CHX) prepared fresh and added to all growth-media, D-PBS, and trypsin at a final concentration of 100 µg/mL. The fresh DMEM was removed from the cells and replaced with DMEM+CHX for 10 min of incubation at 37°C. The cells were then washed with PBS+CHX and treated with Trypsin+CHX for 5 min at 37°C. The trypsinized cells were spun down at 1000rpm for 3 min. The pellet was washed with PBS+CHX and transferred to a pre-weighed 1.5mL microfuge tube. The sample was spun down again at 1000rpm for 3 min and the supernatant was aspirated.

Cell pellets were thawed on ice and resuspended in 1:10 volume of pellet weight: lysis buffer (25mM Tris pH 7.5, 15mM MgCl2, 150mM NaCl, 1% Triton X-100, 1mM DTT, 8% Glycerol, 200U/mL SuperaseIN, 0.2mg/mL CHX, protease inhibitor, and 25U/mL Turbo DNAse). To lyse the pellets, they were vortexed for 30 s and incubated on ice for 30s three times, followed by incubation on ice for 20 min, with vortexing every 5 min. The lysates were then spun at 3700 rpm at 4°C for 5 min. The supernatant was extracted and spun again at 9700 rpm for 5 min. The subsequent supernatant was moved to a fresh tube and used for fractionation. The samples were gently loaded onto the top of a 10-45% sucrose gradient (10% or 45% sucrose, 25mM Tris pH 7.5, 150mM NaCl, 15mM MgCl, 1mM DTT, 1mM CHX, protease inhibitor) prepared in Beckman-Coulter tubes. The sample gradients were then loaded into the pre-chilled SW41 rotor and spun at 40,000 rpm for 2.5 hours at 4 C (Optima XE-90 Ultracentrifuge). The samples were then run through the fractionator (Biocomp Fractionator and Gradient Station) and collected with the fraction collector (Gilson Fraction Collector FC203B). The fractionator was set to use an LED at 260nm, collect 10 data points/mm, separate 580 μL/fraction, and the piston speed was set to 0.2mm/s. The fractions were stored at –20°C following their collection.

### Polysome Profiling Analysis

To correct any shifts in the sucrose gradient position, the number required for the max absorbance (free peak) to reach 0.2 mm was subtracted from all x-axis positions. For absorbance normalization, all absorbance values were divided by the max absorbance. Then, the minimum absorbance was subtracted from all absorbance value to baseline all absorbance values for comparison.

The area under the curve (AUC) between each peak was calculated using integration by trapezoid method using the R package Pracma. The boundaries for each peak were automatically determined by calculating the trough for each peak in R. Disomes were not included as polysomes. Then, the ratio was calculated by dividing the peak AUC to the total AUC.

### AlphaFold3 model and cryo-EM density visualization

The full length PRRC2 sequence was submitted to AlphaFold3 server. The seeds were: PRRC2A 1430183606, PRRC2B 84150108, PRRC2C 629483028. The seed for AlphaFold3 predictions with eIF3 core and PRRC2C residues 506-591 is 592412087.

All models and cryo-EM map (emd3057) were visualized in UCSF ChimeraX(1.1)^125^. The experimental model and cryo-EM map were downloaded from the RCSB Protein Data Bank. The AlphaFold3 model was aligned to the published experimental model (PDB ID:9CPA) in ChimeraX, using matchmaker.

### Western Blot

Cell lysates were collected by directly applying lysis buffer (2% SDS in Tris-HCL pH 7.5) to a 6-well plate at 70-90% confluency. Cell lysates were sonicated and clarified by centrifuging at 15,000g for 5 min. The supernatant was collected and protein concentration was determined using Pierce™ BCA™ Protein Assay (Thermo Fisher). A modified Laemmli sample buffer (with 1xSDS) was added to the sample. Samples were boiled for 5 min at 95°C. Then, equal amounts of protein(15-20 µg) were loaded into each well and resolved with a 10% acrylamide gel. Proteins were transferred onto PVDF membranes using the Trans-Blot Turbo System (BioRad). Membranes were cut and then blocked in either 5% skim milk or 5% BSA (for phosphorylation) in TBST (0.1% Tween-20) for 1-2 hours at room temperature before overnight primary incubation at 4°C. After washing, HRP-conjugated secondaries were incubated for 1 hour at room temperature. Proteins were detected using chemiluminescence substrate. Blot for phospho-eIF2α was stripped using mild stripping buffer (pH 2.2) and used to probe for total eIF2α.

### Gene Ontology Enrichment Analysis

Gene Ontology enrichment analysis was performed in g:Profiler (https://biit.cs.ut.ee/gprofiler/gost)^126^.

### Mass Spectrometry Data Deposition

All proteomics data is available at the Mass Spectrometry Interactive Virtual Environment (MassIVE, http://massive.ucsd.edu/). The total proteome data for PRRC2 KO* cells: MSV000100946 (ftp://massive-ftp.ucsd.edu/v12/MSV000100946/). The BioID data: MSV000100945 (ftp://massive-ftp.ucsd.edu/v12/MSV000100945/). The AP-MS data: MSV000100944 (ftp://massive-ftp.ucsd.edu/v12/MSV000100944/)

## Code Availability

The AlphaFlex analysis scripts can be found in a .ZIP file in the /alphaflex folder in the IDPConformerGenerator GitHub repository.

## ACKNOWLEDGEMENT

The authors wish to thank C. Boone for discussions on genetic interactions, M.S. Choe for helping E.K. use the polysome fractionator, J.Moffat and Y. Xiao for help in CRISPR Cas9-mediated genome engineering. We further wish to thank the members of the core facilities located at Hospital for Sick Children, Toronto, Canada, including L. Baev at SPARC BioCentre, C. Fladd at SPARC Drug Discovery, K. Lau and P. Paroutis at the Imaging Facility for use of their equipment, technical support and training.

## Funding

This work was supported by the Natural Sciences and Engineering Research Council of Canada (NSERC; RGPIN-2022-04849 to J.-Y.Y.; RGPIN-2024-05725 to J.D.F.-K.), Canada Foundation for Innovation (CFI 41430 to J.-Y.Y.; 44381 to O.Z.), National Institutes of Health (NIH R01GM127627 to T Head-Gordon and J.D.F.-K.) and the Canadian Institutes of Health Research (481147 to J.-Y.Y; PJT-148532 to A.M.M and J.D.F.-K.). J.Q.H. is funded by a Canada Graduate Scholarship-Doctoral from NSERC. J.R., J.D.F.-K., and J.-Y.Y. were supported by the Canada Research Chairs program.

## AUTHOR CONTRIBUTIONS

Conceptualization, J.Q.H., J.-Y.Y.; Methodology, J.Q.H., K.J.S., E.K., O.Z.; Software, Z.H.L.; Validation, R.W.C., J.Q.H.; Formal Analysis, J.Q.H.; Investigation, J.Q.H., E.K., Z.H.L.; Resources, K.J.S., T.H.H., S.M.T.A., K.G.; Data Curation, J.Q.H.; Writing – Original Draft, J.Q.H., J.-Y.Y.; Writing – Review & Editing, J.Q.H., J.-Y.Y., O.Z., A.M., J.D.F.-K., J.R.; Visualization, J.Q.H., J.-Y.Y.; Supervision, J.-Y.Y.; Project Administration, J.-Y.Y. and Funding Acquisition, J.-Y.Y., J.D.F.-K., O.Z., A.M.M, T Head-Gordon.

## Supplemental Information

**Figure S1. Molecular and Structural Features of PRRC2 proteins and their orthologs**

**A)** Cytoscape network visualization shows proximal interactions extracted from published BioID data (Youn et al. 2018). PRRC2A, PRRC2B and PRRC2C, indicated in rectangles, are identified as high-confidence interactors with baits indicated in oval nodes, whose localization is indicated by their colour. **B)**Phylogenetic tree of PRRC2 orthologs and analysis of their BAT2 domain, coiled-coil (CC), disorder percentage and number of prion-like domains (PrLD). A check mark indicates the presence of the BAT2 domain. The number of coiled-coils was determined from Marcoil. The percent disorder was calculated from Metapredict disorder scores (see Methods). The number of PrLD was calculated from PLAAC. **C)** Violin plots displaying the distribution of radius of gyration, end-to-end distance, asphericity, and solvent-accessible surface area of 10,000 conformations of PRRC2 proteins. The median and mean are indicated by the red “+” and black dot, respectively.

**Figure S2 PRRC2 localization and condensation dynamics**

**A)** The plots show Manders’ overlap coefficient for fluorescence signal in channel 1 to 2 (left) or channel 2 to 1 (right). The left plot shows the proportion of PRRC2, G3BP1, and UBAP2L signals overlapping with G3BP1. The right measures the proportion of G3BP1 signal overlapping with PRRC2 proteins, G3BP1 and UBAP2L (The letters in parentheses denote G3BP1 monoclonal mouse antibody (M) or polyclonal rabbit antibody (R)). The black circle represents the mean—data from three biological replicates. **B)** The violin plots show the distributions of foci diameter (left) and circularity (right) for G3BP1, PRRC2, and UBAP2L. Wilcoxon rank-sum test comparing G3BP1 foci diameter between SGs induced by the two stressors. **** *P* <0.0001. Data from three biological replicates. **C)** Representative max intensity projections of HAP1 cells endogenously expressing mNeonGreen-PRRC2B and G3BP1-mScarlet (top) and mNeonGreen-PRRC2C and G3BP1-mScarlet (bottom). The white arrow in the inset is pointing to newly formed SGs. Scale bar: 10 µm.

**Figure S3 PRRC2A and PRRC2B BioID Biotinylation and Localization**

**A)** Western blot of BioID cell lysates, probed for biotinylated proteins using Streptavidin-HRP. Cleared lysates after cell lysis in modified RIPA buffer were used. **B)** The heatmap shows the Jaccard similarity indices for high-confidence interactors identified in AP-MS and BioID experiments. **C)** Representative single-plane immunofluorescence images of HEK293 Flp-In TREx cells expressing PRRC2 fused to MiniTurbo-FLAG after 35 min of NaAsO_2_ stress and 30 min of biotin labelling. TIAR was used as a stress granule marker. Streptavidin staining shows the localization of biotinylated proteins. Scale bar: 10 µm

**Figure S4 Stress-Induced Rearrangements of Protein Interaction Network for PRRC2A and PRRC2B**

**A)** Venn Diagrams show the overlap in SG tier 1 proteins identified as high-confidence interactors between PRRC2A and PRRC2B across different conditions. **B)** The line plot shows the abundance changes for components of the eIF3 complex identified as high-confidence interactors by PRRC2B under no stress and stress conditions. Preys whose abundance changes by ≥ 1.5-fold are indicated in black with orange lines, and prey decreasing by ≥ 1.5-fold are indicated in blue with blue lines. **C)** The line plot shows the abundance changes for other translation initiation factors (eIF proteins) identified as high-confidence interactors by PRRC2A or PRRC2B under no stress or stress conditions. Preys whose abundance changes by ≥ 1.5-fold are indicated in black with orange lines, and preys decreasing by ≥ 1.5-fold are indicated in blue with blue lines. **D)** The line plot shows the abundance changes for ribosome subunits identified as high-confidence interactors by PRRC2A or PRRC2B under no stress or stress conditions. Preys whose abundance changes by ≥ 1.5-fold are indicated in black with orange lines, and preys decreasing by ≥ 1.5-fold are indicated in blue with blue lines. **E)** The line plot shows the abundance changes for 5’ mRNA decay pathway components identified as high-confidence interactors by PRRC2A or PRRC2B across no stress or stress conditions. Preys whose abundance changes by ≥ 1.5-fold are indicated in black with orange lines, and preys decreasing by ≥ 1.5-fold are indicated in blue with blue lines.

**Figure S5 PRRC2B Proximal Interactions with Proteins Involved in DNA Damage and Cell Cycle**

**A)** Dot plot of nuclear importers proximally associated with PRRC2 proteins. **B)** Schematic of putative Nuclear Localization Signals in PRRC2 proteins. NLS sequences were found using the ELM database. PY-NLS was identified using regular expression (see Methods).**C)** Representative immunofluorescence max intensity projections showing localization of endogenous PRRC2B in HeLa Flp-In TREx cells. Scale bar: 10 µm. **D)** Line plot showing log_2_(Prey Intensity) changes in different conditions for PRRC2B nuclear preys. Preys increasing ≥1.5 fold are coloured in orange. Preys decreasing ≥1.5 fold are coloured in blue. **E)** Line plot showing log_2_(Prey Intensity) changes in different conditions for DNA damage proteins. Preys increasing ≥1.5 fold are coloured in orange. Preys decreasing ≥1.5 fold are coloured in blue. **F)** Dot plot highlighting DNA damage prey. **G)** PRRC2B cell cycle preys. Line plot showing log_2_(Prey Intensity) changes in different conditions for cell cycle proteins. Preys increasing ≥1.5 fold are coloured in orange. Preys decreasing ≥1.5 fold are coloured in blue.**H)** Dot plot highlighting cell cycle-associated preys. **I)** Evolutionary signatures of degron SLiMs and their location in human PRRC2 proteins. DEG_APCC_KENBOX_2 is a destruction motif targeted by the Anaphase-promoting ubiquitin ligase complex.

**Figure S6 Proteomics Quality Control**

**A)** Schematic for whole proteome analysis. Cells were collected from a 12-well plate. Peptides were extracted using S-trap Micro. Samples were analyzed using DIA-MS and processed in DIA-NN. MSStats was used for downstream analysis. **B)** Violin plots showing the coefficient of variation between replicates at the fragment level. Dashed line = 15% variation. **C)** Hierarchical clustering of protein abundances in each replicate. The dendrogram was generated using Euclidean distance and Ward’s linkage. **D)** Peptide abundance log_2_(Fold-Change) across *PRRC2A* mutants. The dashed line indicates the CRISPR/Cas9 target site. Fragment intensities were summed into peptide-level information for calculating Fold-Change. **E)** Peptide abundance log_2_(Fold-Change) across PRRC2B in *PRRC2B* and *PRRC2C* double knockouts. The dashed line indicates the CRISPR/Cas9 target site. Fragment intensities were summed into peptide-level information for calculating Fold-Change.

**Figure S7 PRRC2 single KO* clones SG Assembly Phenotype**

**A)** Representative immunofluorescence max intensity projections of G3BP1 in *PRRC2* single mutants. Scale bar: 20 µm. **B)** Violin plot showing cell area (µm^2^) of *PRRC2* single mutants. Wilcox test with Holm correction, mutants compared to WT. * P-adjusted < 0.05, ** P< 0.01, *** P <0.001.**C)** Violin plots showing the number of G3BP1 foci in single mutants. Statistics were performed with the Wilcox test with Holm correction, mutants compared to WT. * P-adjusted < 0.05, ** P< 0.01, *** P <0.001. The data show combined results from 3 biological replicates. n = total number of cells **D)** Scatter plot of G3BP1 foci number and cell area for wildtype cells in oxidative stress and hyperosmotic stress. **E)** Western blots showing G3BP1, phospho-eIF2α and total eIF2α in *PRRC2* single mutants in no stress, NaAsO_2_ stress (left) and NaCl stress (right).

**Figure S8 ABC KO* clones SG Assembly and PB Assembly Phenotype**

**A)** Representative immunofluorescence max intensity projections of G3BP1 and DCP1A in *ABC* KO* clones. Scale bar: 10 µm. **B)** Western blot showing G3BP1, phospho-eIF2α and total eIF2α in *ABC* KO* clones. **C)** Violin plot showing cell area (µm^2^) of *ABC* KO* clones. Statistics were performed with the Wilcox test with Holm correction, mutants compared to WT. * P-adjusted < 0.05, ** P< 0.01, *** P <0.001. **D)** Violin plots showing the number of DCP1A for *ABC* KO* clones. Statistics were performed with the Wilcox test with Holm correction, mutants compared to WT. * P-adjusted < 0.05, ** P< 0.01, *** P <0.001. The data show combined results from 3 biological replicates. n = total number of cells. **E)** Scatter plot of DCP1A foci number and cell area for wildtype cells in oxidative stress and hyperosmotic stress.

**Figure S9 PRRC2 Protein DepMap Network**

**A)** Cytoscape network showing *PRRC2* genes and their codependent genes from DepMap. Only genes with |correlation| ≥ 0.15 are displayed. **B)** Violin plot showing log_2_ fold-change for protein abundance in *ABC* KO* clones. Proteins were categorized into eIF3d translation targets and non-eIF3d targets. Statistics were performed with the Wilcoxon rank-sum test. **** p-value < 0.0001. **C)** Violin plot showing log_2_ fold-change for protein abundance in *ABC* KO* clones. Proteins were categorized into eIF4G2 translation targets and non-eIF4G2 targets. Statistics were performed with the Wilcoxon rank-sum test. **** p-value < 0.0001. **D)** Violin plot showing log_2_ fold-change for protein abundance in *ABC* KO* clones. Proteins were categorized into groups based on the presence or absence of upstream open reading frames in their mRNA. Statistics were performed with the Wilcoxon rank-sum test. **** p-value < 0.0001. **E)** Violin plot showing log_2_ fold-change for protein abundance in *ABC* KO* clones. Proteins were categorized into groups based on the presence or absence of overlapping open reading frames in their mRNA. Statistics were performed with the Wilcoxon rank-sum test. Ns, not significant, ** p-value < 0.01, *** p-value < 0.001

**Figure S10 PRRC2C AP-MS and SLiM Analysis A)** Barplot showing the number of high-confidence prey detected for PRRC2C full-length and deletion mutants. **B)** Barplot showing the log_2_ (intensity) of PRRC2C in full-length and deletion mutants. **C)** Selected GO terms to highlight alterations in PRRC2C AP-MS interactome. ns, not significant. **D)** Density plot showing the distribution of solvent-accessible surface area for TRG_NLS_MonoCore_2 in AlphaFlex models of PRRC2C full length and PRRC2C-ΔBAT2. This is an NLS SLiM. 10,000 conformations were analyzed. **E)** Density plot showing the distribution of solvent-accessible surface area for LIG_CtBP_PxDLS_1 in AlphaFlex models of PRRC2C full length and PRRC2-ΔBAT2. This is a binding SLiM for CtBP corepressors. 10,000 conformations were analyzed. **F)** Bar plot showing the number of high confidence preys (AvgP ≥ 0.98) with ≥1.5-fold change in PRRC2C deletion mutant compared to full-length PRRC2C. **G)** Line plot highlighting the 5 preys that showed a 1.5-fold decrease in RG isoform deletion. **H)** AlphaFold3 predicts the presence of α helices in PRRC2 proteins. The regions predicted to form coiled-coils (≥99% confidence) are coloured in bright green and are indicated by the amino acid number.

**Table S1**. Evolutionary Signatures and NARDINI+ scores for PRRC2 proteins

**Table S2**. AP-MS tables for PRRC2 proteins

**Table S3**. GO enrichment tables for PRRC2 AP-MS

**Table S4**. BioID tables for PRRC2 proteins

**Table S5**. GO enrichment tables for PRRC2A and PRRC2B BioID

**Table S6**. Genotyping Results for PRRC2 CRISPR-Cas9 mutants

**Table S7**. MSstats protein comparisons

**Table S8**. DepMap Gene Co-essentially table

**Table S9**. GO enrichment tables for PRRC2C deletion construct AP-MS

## Notes

### Competing Interest Statement

The authors have declared no competing interest.

## REFERENCES

1. de la Parra, C., Walters, B. A., Geter, P. & Schneider, R. J. Translation initiation factors and their relevance in cancer. Curr. Opin. Genet. Dev. 48, 82–88 (2018).

2. Kapur, M. & Ackerman, S. L. mRNA Translation Gone Awry: Translation Fidelity and Neurological Disease. Trends Genet. TIG 34, 218–231 (2018).

3. Sonenberg, N. & Hinnebusch, A. G. Regulation of translation initiation in eukaryotes: mechanisms and biological targets. Cell 136, 731–745 (2009).

4. Vaklavas, C., Blume, S. W. & Grizzle, W. E. Translational Dysregulation in Cancer: Molecular Insights and Potential Clinical Applications in Biomarker Development. Front. Oncol. 7, 158 (2017).

5. Hofmann, S., Kedersha, N., Anderson, P. & Ivanov, P. Molecular mechanisms of stress granule assembly and disassembly. Biochim. Biophys. Acta - Mol. Cell Res. 1868, 118876 (2021).

6. Ivanov, P., Kedersha, N. & Anderson, P. Stress granules and processing bodies in translational control. Cold Spring Harb. Perspect. Biol. 11, (2019).

7. Costa-Mattioli, M. & Walter, P. The integrated stress response: From mechanism to disease. Science 368, (2020).

8. Pakos-Zebrucka, K. et al. The integrated stress response. EMBO Rep. 17, 1374–1395 (2016).

9. Protter, D. S. W. & Parker, R. Principles and Properties of Stress Granules. Trends Cell Biol. 26, 668–679 (2016).

10. Anderson, P. & Kedersha, N. Stress granules: the Tao of RNA triage. Trends Biochem. Sci. 33, 141–150 (2008).

11. Wolozin, B. & Ivanov, P. Stress granules and neurodegeneration. Nat. Rev. Neurosci. 20, 649–666 (2019).

12. Yang, P. et al. G3BP1 Is a Tunable Switch that Triggers Phase Separation to Assemble Stress Granules. Cell 181, 325–345.e28 (2020).

13. Sanders, D. W. et al. Competing Protein-RNA Interaction Networks Control Multiphase Intracellular Organization. Cell 181, 306–324.e28 (2020).

14. Mittag, T. & Parker, R. Multiple Modes of Protein–Protein Interactions Promote RNP Granule Assembly. J. Mol. Biol. 430, 4636–4649 (2018).

15. Banani, S. F. et al. Compositional Control of Phase-Separated Cellular Bodies. Cell 166, 651–663 (2016).

16. Smith, J. & Bartel, D. P. The G3BP stress-granule proteins reinforce the integrated stress response translation programme. Nat. Cell Biol. 28, 135–148 (2026).

17. Youn, J. Y. et al. High-Density Proximity Mapping Reveals the Subcellular Organization of mRNA-Associated Granules and Bodies. Mol. Cell 69, 517–532.e11 (2018).

18. Markmiller, S. et al. Context-Dependent and Disease-Specific Diversity in Protein Interactions within Stress Granules. Cell 172, 590–604.e13 (2018).

19. Laver, J. D. et al. The RNA-Binding Protein Rasputin/G3BP Enhances the Stability and Translation of Its Target mRNAs. Cell Rep. 30, 3353–3367.e7 (2020).

20. Meyer, C., Garzia, A., Morozov, P., Molina, H. & Tuschl, T. The G3BP1-Family-USP10 Deubiquitinase Complex Rescues Ubiquitinated 40S Subunits of Ribosomes Stalled in Translation from Lysosomal Degradation. Mol. Cell 77, 1193–1205.e5 (2020).

21. Luo, E.-C. et al. Large-scale tethered function assays identify factors that regulate mRNA stability and translation. Nat. Struct. Mol. Biol. 27, 989–1000 (2020).

22. Khandjian, E. W., Corbin, F., Woerly, S. & Rousseau, F. The fragile X mental retardation protein is associated with ribosomes. Nat. Genet. 12, 91–93 (1996).

23. Stefani, G., Fraser, C. E., Darnell, J. C. & Darnell, R. B. Fragile X mental retardation protein is associated with translating polyribosomes in neuronal cells. J. Neurosci. Off. J. Soc. Neurosci. 24, 7272–7276 (2004).

24. Feng, Y. et al. FMRP associates with polyribosomes as an mRNP, and the I304N mutation of severe fragile X syndrome abolishes this association. Mol. Cell 1, 109–118 (1997).

25. Thomas, M. G., Martinez Tosar, L. J., Desbats, M. A., Leishman, C. C. & Boccaccio, G. L. Mammalian Staufen 1 is recruited to stress granules and impairs their assembly. J. Cell Sci. 122, 563–573 (2009).

26. Dugré-Brisson, S. et al. Interaction of Staufen1 with the 5’ end of mRNA facilitates translation of these RNAs. Nucleic Acids Res. 33, 4797–4812 (2005).

27. Wu, R. et al. A novel m6A reader Prrc2a controls oligodendroglial specification and myelination. Cell Res. 29, 23–41 (2019).

28. Groza, T. et al. The International Mouse Phenotyping Consortium: comprehensive knockout phenotyping underpinning the study of human disease. Nucleic Acids Res. 51, D1038–D1045 (2023).

29. Bohlen, J., Roiuk, M., Neff, M. & Teleman, A. A. PRRC2 proteins impact translation initiation by promoting leaky scanning. Nucleic Acids Res. 51, 3391–3409 (2023).

30. Zhang, T. et al. RNA-binding protein Nocte regulates Drosophila development by promoting translation reinitiation on mRNAs with long upstream open reading frames. Nucleic Acids Res. 52, 885–905 (2024).

31. Jiang, F. et al. RNA binding protein PRRC2B mediates translation of specific mRNAs and regulates cell cycle progression. Nucleic Acids Res. gkad322 (2023) doi:10.1093/nar/gkad322.

32. Goldberg, N. et al. The RNA-binding protein PRRC2B preserves 5′ TOP mRNA during starvation to maintain ribosome biogenesis during nutrient recovery. Nucleic Acids Res. 53, gkaf1334 (2025).

33. Panhale, A. et al. CAPRI enables comparison of evolutionarily conserved RNA interacting regions. Nat. Commun. 10, (2019).

34. Bai, G. et al. The molecular characteristics in different procedures of spermatogenesis. Gene 826, 146405 (2022).

35. Tan, X., Zheng, C., Zhuang, Y., Jin, P. & Wang, F. The m6A reader PRRC2A is essential for meiosis I completion during spermatogenesis. Nat. Commun. 14, 1636 (2023).

36. Li, S. et al. The Role of PRRC2B in Cerebral Vascular Remodeling Under Acute Hypoxia in Mice. Adv. Sci. https://doi.org/10.1002/advs.202300892 (2023) doi:10.1002/advs.202300892.

37. Zhang, Y. et al. PRRC2B modulates oligodendrocyte progenitor cell development and myelination by stabilizing Sox2 mRNA. Cell Rep. 43, 113930 (2024).

38. Li, X. et al. Microglial Prrc2a regulates microglia-neuron interaction in cerebellum and motor functions in mice. iScience 28, 113949 (2025).

39. Shi, H., Wei, J. & He, C. Where, When, and How: Context-Dependent Functions of RNA Methylation Writers, Readers, and Erasers. Mol. Cell 74, 640–650 (2019).

40. Zaccara, S., Ries, R. J. & Jaffrey, S. R. Reading, writing and erasing mRNA methylation. Nat. Rev. Mol. Cell Biol. 20, 608–624 (2019).

41. Jackson, R. J., Hellen, C. U. T. & Pestova, T. V. The mechanism of eukaryotic translation initiation and principles of its regulation. Nat. Rev. Mol. Cell Biol. 11, 113–127 (2010).

42. Wolf, D. A., Lin, Y., Duan, H. & Cheng, Y. EIF-Three to Tango: Emerging functions of translation initiation factor eIF3 in protein synthesis and disease. J. Mol. Cell Biol. 12, 403–409 (2020).

43. Valášek, L. S. et al. Embraced by eIF3: structural and functional insights into the roles of eIF3 across the translation cycle. Nucleic Acids Res. 45, 10948–10968 (2017).

44. Ide, N. A., Noorwez, S., Xu, C. & Aitken, C. E. eIF3: a critical player in mRNA recruitment to the ribosome with emerging roles across translation. Biochem. Soc. Trans. 53, 1527–1541 (2025).

45. Roy, B. et al. The h subunit of eIF3 promotes reinitiation competence during translation of mRNAs harboring upstream open reading frames. RNA 16, 748–761 (2010).

46. Mohammad, M. P., Munzarová Pondelícková, V., Zeman, J., Gunišová, S. & Valášek, L. S. In vivo evidence that eIF3 stays bound to ribosomes elongating and terminating on short upstream ORFs to promote reinitiation. Nucleic Acids Res. 45, 2658–2674 (2017).

47. Munzarová, V. et al. Translation reinitiation relies on the interaction between eIF3a/TIF32 and progressively folded cis-acting mRNA elements preceding short uORFs. PLoS Genet. 7, e1002137 (2011).

48. Lambert, J. P., Tucholska, M., Go, C., Knight, J. D. R. & Gingras, A. C. Proximity biotinylation and affinity purification are complementary approaches for the interactome mapping of chromatin-associated protein complexes. J. Proteomics 118, 81–94 (2015).

49. Gingras, A. C., Abe, K. T. & Raught, B. Getting to know the neighborhood: using proximity-dependent biotinylation to characterize protein complexes and map organelles. Curr. Opin. Chem. Biol. 48, 44–54 (2019).

50. Branon, T. C. et al. Efficient proximity labeling in living cells and organisms with TurboID. Nat. Biotechnol. 36, 880–887 (2018).

51. Youn, J. Y. et al. Properties of Stress Granule and P-Body Proteomes. Mol. Cell 76, 286–294 (2019).

52. Blum, M. et al. InterPro: the protein sequence classification resource in 2025. Nucleic Acids Res. 53, D444–D456 (2025).

53. Liu, Z. H., et al. AlphaFlex: Ensembles of the human proteome representing disordered regions. bioRxiv 2025.11.24.690279 (2025) doi:10.1101/2025.11.24.690279.

54. Sabari, B. R., Hyman, A. A. & Hnisz, D. Functional specificity in biomolecular condensates revealed by genetic complementation. Nat. Rev. Genet. 26, 279–290 (2025).

55. Zarin, T. et al. Proteome-wide signatures of function in highly diverged intrinsically disordered regions. eLife 8, 1–26 (2019).

56. Zarin, T. et al. Identifying molecular features that are associated with biological function of intrinsically disordered protein regions. eLife 10, 1–23 (2021).

57. Ruff, K. M. et al. Molecular grammars of predicted intrinsically disordered regions that span the human proteome. Cell 0, (2025).

58. Pei, G., Lyons, H., Li, P. & Sabari, B. R. Transcription regulation by biomolecular condensates. Nat. Rev. Mol. Cell Biol. 26, 213–236 (2025).

59. Ozdilek, B. A. et al. Intrinsically disordered RGG/RG domains mediate degenerate specificity in RNA binding. Nucleic Acids Res. 45, 7984–7996 (2017).

60. Thandapani, P., O’Connor, T. R., Bailey, T. L. & Richard, S. Defining the RGG/RG Motif. Mol. Cell 50, 613–623 (2013).

61. Chong, P. A., Vernon, R. M. & Forman-Kay, J. D. RGG/RG Motif Regions in RNA Binding and Phase Separation. J. Mol. Biol. 430, 4650–4665 (2018).

62. Chowdhury, M. N. & Jin, H. The RGG motif proteins: Interactions, functions, and regulations. Wiley Interdiscip. Rev. RNA 14, e1748 (2023).

63. Pritišanac, I. et al. A Functional Map of the Human Intrinsically Disordered Proteome. 2024.03.15.585291 Preprint at 10.1101/2024.03.15.585291 (2024).

64. Aulas, A. et al. G3BP1 promotes stress-induced RNA granule interactions to preserve polyadenylated mRNA. J. Cell Biol. 209, 73–84 (2015).

65. Jain, S. et al. ATPase modulated stress granules contain a diverse proteome and substructure. Cell 164, 487 (2016).

66. Guillén-Boixet, J. et al. RNA-Induced Conformational Switching and Clustering of G3BP Drive Stress Granule Assembly by Condensation. Cell 181, 346–361.e17 (2020).

67. Lehner, B. et al. Analysis of a high-throughput yeast two-hybrid system and its use to predict the function of intracellular proteins encoded within the human MHC class III region. Genomics 83, 153–167 (2004).

68. Demichev, V., Messner, C. B., Vernardis, S. I., Lilley, K. S. & Ralser, M. DIA-NN: Neural networks and interference correction enable deep proteome coverage in high throughput. Nat. Methods 17, 41–44 (2020).

69. Teo, G. et al. SAINTq: Scoring protein-protein interactions in affinity purification – mass spectrometry experiments with fragment or peptide intensity data. Proteomics 16, 2238–2245 (2016).

70. Tsai, Y.-C. et al. Dynamic Proximal Interactomics and Chemical Genetic Screening Reveal CCR4-NOT Sequestration in Stress Granules as a Mechanism for Transcript Stabilization. 2025.02.04.636058 Preprint at 10.1101/2025.02.04.636058 (2025).

71. Piette, B. L. et al. Comprehensive interactome profiling of the human Hsp70 network highlights functional differentiation of J domains. Mol. Cell 81, 2549–2565.e8 (2021).

72. Riggs, C. L. et al. UBAP2L contributes to formation of P-bodies and modulates their association with stress granules. J. Cell Biol. 223, e202307146 (2024).

73. Meyer, K. D. et al. 5′ UTR m6A Promotes Cap-Independent Translation. Cell 163, 999–1010 (2015).

74. Mukhopadhyay, S., Amodeo, M. E. & Lee, A. S. Y. eIF3d controls the persistent integrated stress response. Mol. Cell 83, 3303–3313.e6 (2023).

75. Wu, C. C.-C., Peterson, A., Zinshteyn, B., Regot, S. & Green, R. Ribosome Collisions Trigger General Stress Responses to Regulate Cell Fate. Cell 182, 404–416.e14 (2020).

76. D’Orazio, K. N. & Green, R. Ribosome states signal RNA quality control. Mol. Cell 81, 1372–1383 (2021).

77. An, H., Tan, J. T. & Shelkovnikova, T. A. Stress granules regulate stress-induced paraspeckle assembly. J. Cell Biol. 218, 4127–4140 (2019).

78. Róna, G. et al. NLS copy-number variation governs efficiency of nuclear import--case study on dUTPases. FEBS J. 281, 5463–5478 (2014).

79. Sahin, U. et al. Oxidative stress-induced assembly of PML nuclear bodies controls sumoylation of partner proteins. J. Cell Biol. 204, 931–945 (2014).

80. Dyakov, B. J. A. et al. Spatial proteomic mapping of human nuclear bodies reveals new functional insights into RNA regulation. 2024.07.03.601239 Preprint at 10.1101/2024.07.03.601239 (2024).

81. Lallemand-Breitenbach, V. & de Thé, H. PML nuclear bodies. Cold Spring Harb. Perspect. Biol. 2, a000661 (2010).

82. Cooke, M. S., Evans, M. D., Dizdaroglu, M. & Lunec, J. Oxidative DNA damage: mechanisms, mutation, and disease. FASEB J. Off. Publ. Fed. Am. Soc. Exp. Biol. 17, 1195–1214 (2003).

83. Chang, H. R. et al. The functional roles of PML nuclear bodies in genome maintenance. Mutat. Res. Mol. Mech. Mutagen. 809, 99–107 (2018).

84. Alamillo, L. et al. Deuterium labeling enables proteome-wide turnover kinetics analysis in cell culture. *Cell Rep*. Methods 5, 101104 (2025).

85. Smits, A. H. et al. Biological plasticity rescues target activity in CRISPR knock outs. Nat. Methods 16, 1087–1093 (2019).

86. Hubstenberger, A. et al. P-Body Purification Reveals the Condensation of Repressed mRNA Regulons. Mol. Cell 68, 144–157.e5 (2017).

87. Uhlen, M. et al. A pathology atlas of the human cancer transcriptome. Science 357, eaan2507 (2017).

88. Arafeh, R., Shibue, T., Dempster, J. M., Hahn, W. C. & Vazquez, F. The present and future of the Cancer Dependency Map. Nat. Rev. Cancer 25, 59–73 (2025).

89. Smirnova, V. V. et al. Ribosomal leaky scanning through a translated uORF requires eIF4G2. Nucleic Acids Res. 50, 1111–1127 (2022).

90. Billmann, M. et al. A global genetic interaction map of a human cell reveals conserved principles of genetic networks. 2025.06.30.662193 Preprint at 10.1101/2025.06.30.662193 (2025).

91. Pla-Prats, C. & Thomä, N. H. Quality control of protein complex assembly by the ubiquitin-proteasome system. Trends Cell Biol. 32, 696–706 (2022).

92. Padovani, C., Jevtić, P. & Rapé, M. Quality control of protein complex composition. Mol. Cell 82, 1439–1450 (2022).

93. Herrmannová, A. et al. Perturbations in eIF3 subunit stoichiometry alter expression of ribosomal proteins and key components of the MAPK signaling pathways. eLife 13, RP95846 (2024).

94. Weber, R. et al. DAP5 enables main ORF translation on mRNAs with structured and uORF-containing 5’ leaders. Nat. Commun. 13, 7510 (2022).

95. Shestakova, E. D. et al. The Roles of eIF4G2 in Leaky Scanning and Reinitiation on the Human Dual-Coding POLG mRNA. Int. J. Mol. Sci. 24, 17149 (2023).

96. Chen, J. et al. Pervasive functional translation of noncanonical human open reading frames. Science 367, 1140–1146 (2020).

97. Abramson, J. et al. Accurate structure prediction of biomolecular interactions with AlphaFold 3. Nature 630, 493–500 (2024).

98. des Georges, A. et al. Structure of mammalian eIF3 in the context of the 43S preinitiation complex. Nature 525, 491–495 (2015).

99. Cui, D., Pestova, T., Hellen, C. & des Georges, A. The translation initiation factor DHX29 appears to pull on mRNA in a direction opposite to scanning. BioRxiv Prepr. Serv. Biol. 2025.07.13.664561 (2025) doi:10.1101/2025.07.13.664561.

100. Mateju, D. et al. Single-Molecule Imaging Reveals Translation of mRNAs Localized to Stress Granules. Cell 183, 1801–1812.e13 (2020).

101. Wu, X. et al. m6A Reader PRRC2A Promotes Colorectal Cancer Progression via CK1ε-Mediated Activation of WNT and YAP Signaling Pathways. Adv. Sci. 12, 2406935 (2024).

102. Zhou, J. et al. Dynamic m(6)A mRNA methylation directs translational control of heat shock response. Nature 526, 591–594 (2015).

103. Li, Y. et al. Proteogenomic data and resources for pan-cancer analysis. Cancer Cell 41, 1397–1406 (2023).

104. Meril, S. et al. Loss-of-function cancer-linked mutations in the EIF4G2 non-canonical translation initiation factor. Life Sci. Alliance 7, e202302338 (2023).

105. Xiao, Y.-X., Wei, J. & Moffat, J. Protocol for CRISPR-based endogenous protein tagging in mammalian cells. STAR Protoc. 5, 103404 (2024).

106. Conant, D. et al. Inference of CRISPR Edits from Sanger Trace Data. CRISPR J. 5, 123–130 (2022).

107. Emenecker, R. J., Griffith, D. & Holehouse, A. S. Metapredict: a fast, accurate, and easy-to-use predictor of consensus disorder and structure. Biophys. J. 120, 4312–4319 (2021).

108. Lotthammer, J. M. et al. Metapredict enables accurate disorder prediction across the Tree of Life. bioRxiv 10.1101/2024.11.05.622168 (2024)

109. Delorenzi, M. & Speed, T. An HMM model for coiled-coil domains and a comparison with PSSM-based predictions. Bioinformatics 18, 617–625 (2002).

110. Zimmermann, L. et al. A Completely Reimplemented MPI Bioinformatics Toolkit with a New HHpred Server at its Core. J. Mol. Biol. 430, 2237–2243 (2018).

111. Lancaster, A. K., Nutter-Upham, A., Lindquist, S. & King, O. D. PLAAC: A web and command-line application to identify proteins with prion-like amino acid composition. Bioinformatics 30, 2501–2502 (2014).

112. Kumar, M. et al. ELM-the Eukaryotic Linear Motif resource-2024 update. Nucleic Acids Res. 52, D442–D455 (2024).

113. de Castro, E. et al. ScanProsite: detection of PROSITE signature matches and ProRule-associated functional and structural residues in proteins. Nucleic Acids Res. 34, W362–365 (2006).

114. Lee, B. J. et al. Rules for Nuclear Localization Sequence Recognition by Karyopherinβ2. Cell 126, 543–558 (2006).

115. Dyer, S. C. et al. Ensembl 2025. Nucleic Acids Res. 53, D948–D957 (2025).

116. Katoh, K. & Standley, D. M. MAFFT Multiple Sequence Alignment Software Version 7: Improvements in Performance and Usability. Mol. Biol. Evol. 30, 772–780 (2013).

117. Teixeira, J. M. C. et al. IDPConformerGenerator: A Flexible Software Suite for Sampling the Conformational Space of Disordered Protein States. J. Phys. Chem. A 126, 5985–6003 (2022).

118. Liu, Z. H. et al. Local Disordered Region Sampling (LDRS) for ensemble modeling of proteins with experimentally undetermined or low confidence prediction segments. Bioinformatics 39, btad739 (2023).

119. Berman, H. M. et al. The Protein Data Bank. Nucleic Acids Res. 28, 235–242 (2000).

120. Mitternacht, S. FreeSASA: An open source C library for solvent accessible surface area calculations. F1000Research 5, 189 (2016).

121. Tien, M. Z., Meyer, A. G., Sydykova, D. K., Spielman, S. J. & Wilke, C. O. Maximum Allowed Solvent Accessibilites of Residues in Proteins. PLoS ONE 8, e80635 (2013).

122. Shannon, P. et al. Cytoscape: A Software Environment for Integrated Models of Biomolecular Interaction Networks. Genome Res. 13, 2498–2504 (2003).

123. Stirling, D. R. et al. CellProfiler 4: improvements in speed, utility and usability. BMC Bioinformatics 22, 433 (2021).

124. Knight, J. D. R. et al. ProHits-viz: a suite of web tools for visualizing interaction proteomics data. Nat. Methods 2017 147 14, 645–646 (2017).

125. Meng, E. C. et al. UCSF ChimeraX: Tools for structure building and analysis. Protein Sci. Publ. Protein Soc. 32, e4792 (2023).

126. Raudvere, U. et al. g:Profiler: a web server for functional enrichment analysis and conversions of gene lists (2019 update). Nucleic Acids Res. https://doi.org/10.1093/nar/gkz369 (2019) doi:10.1093/nar/gkz369.

