## Supplementary figures and images for "PRRC2A, PRRC2B and PRRC2C are Stress Granule Proteins that Promote Translation Through Association with the eIF3 complex"

### Supplementary Figures

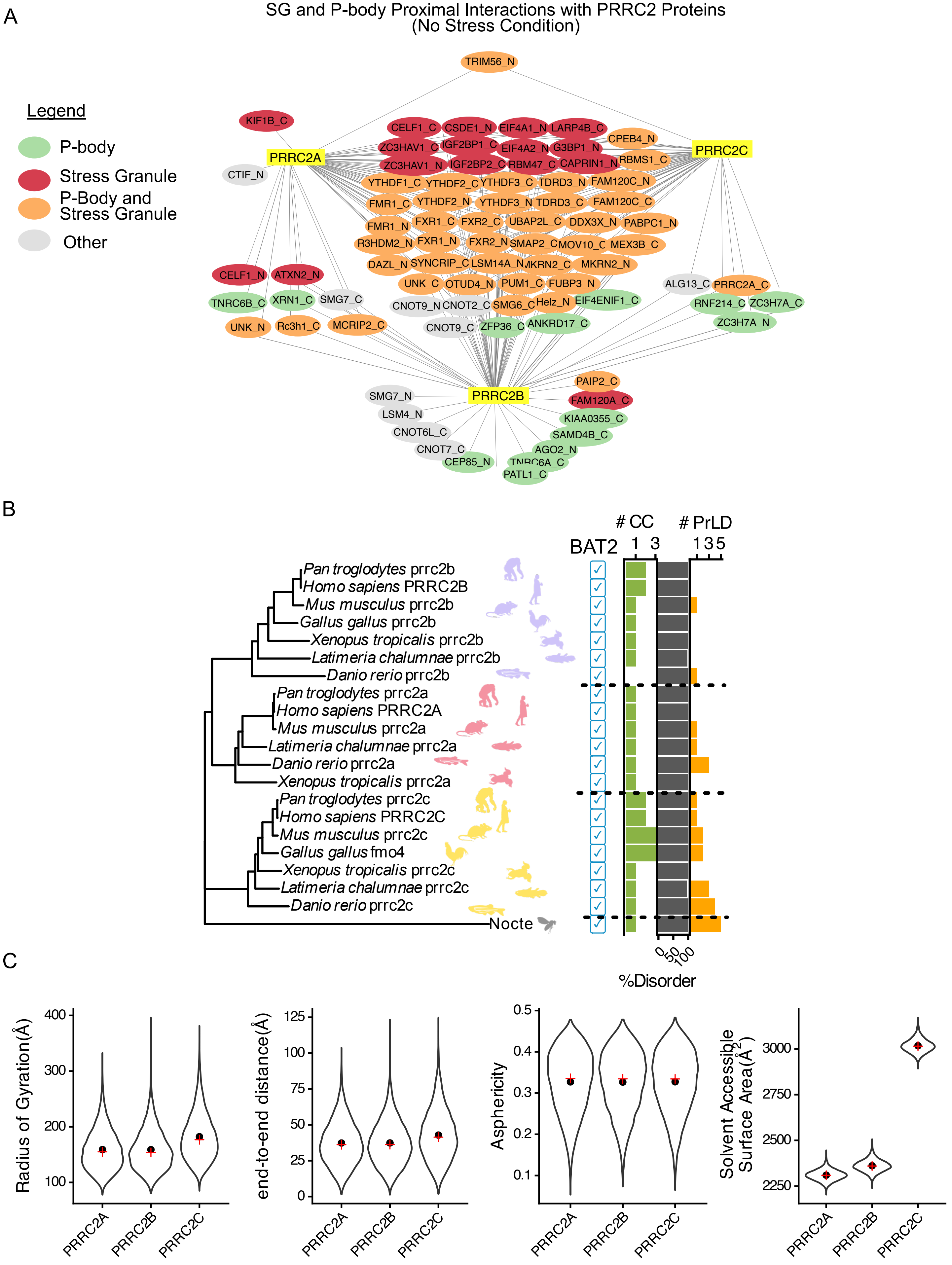

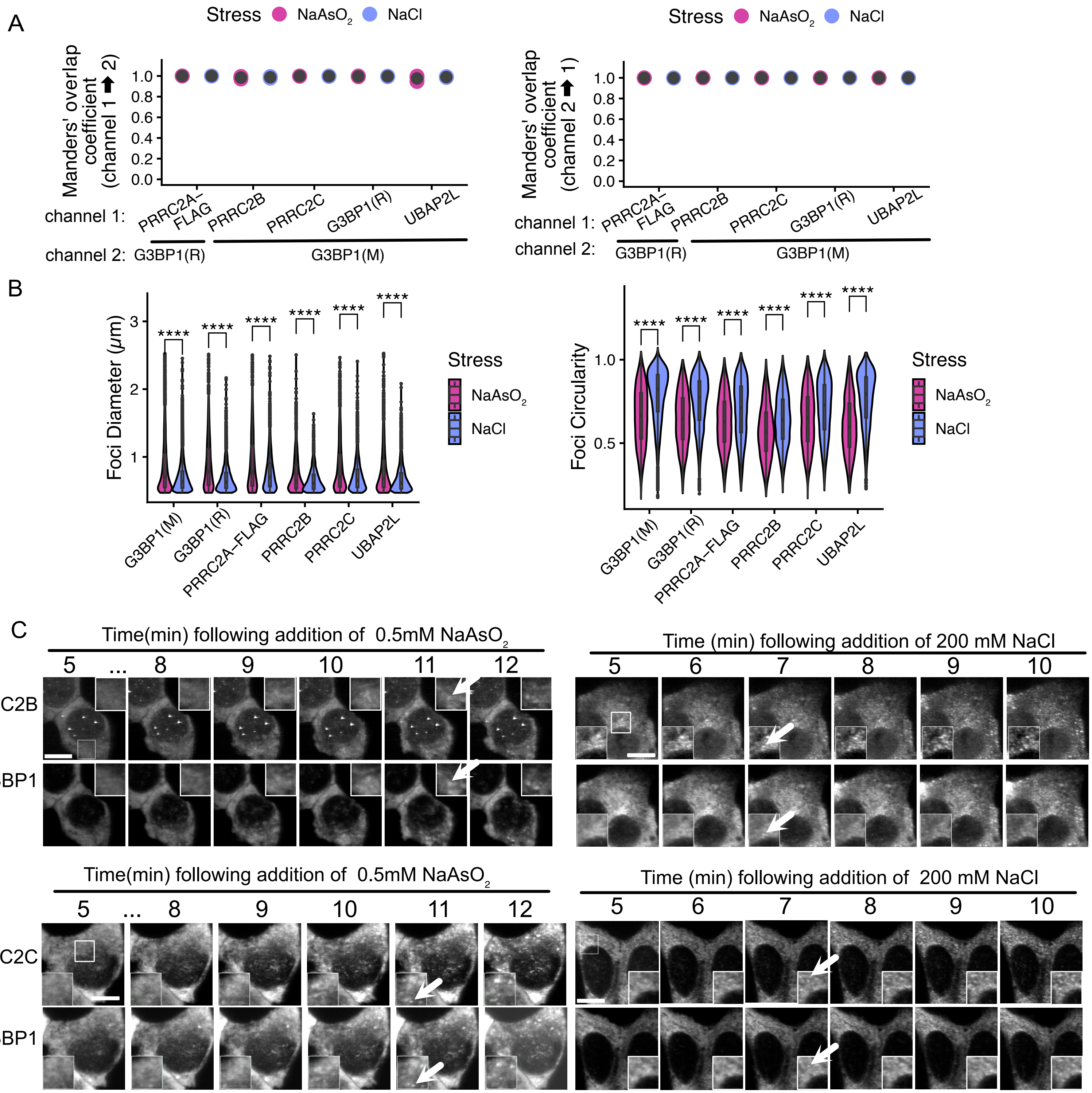

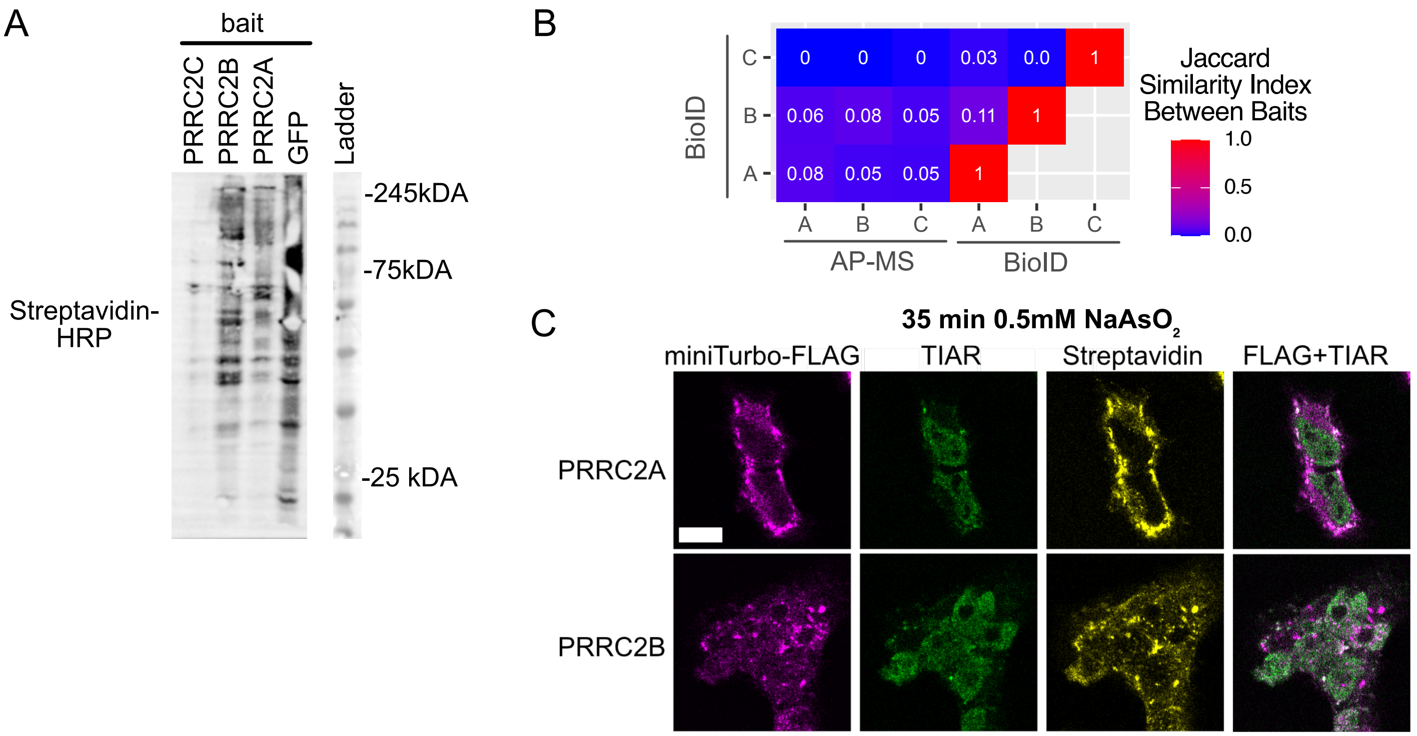

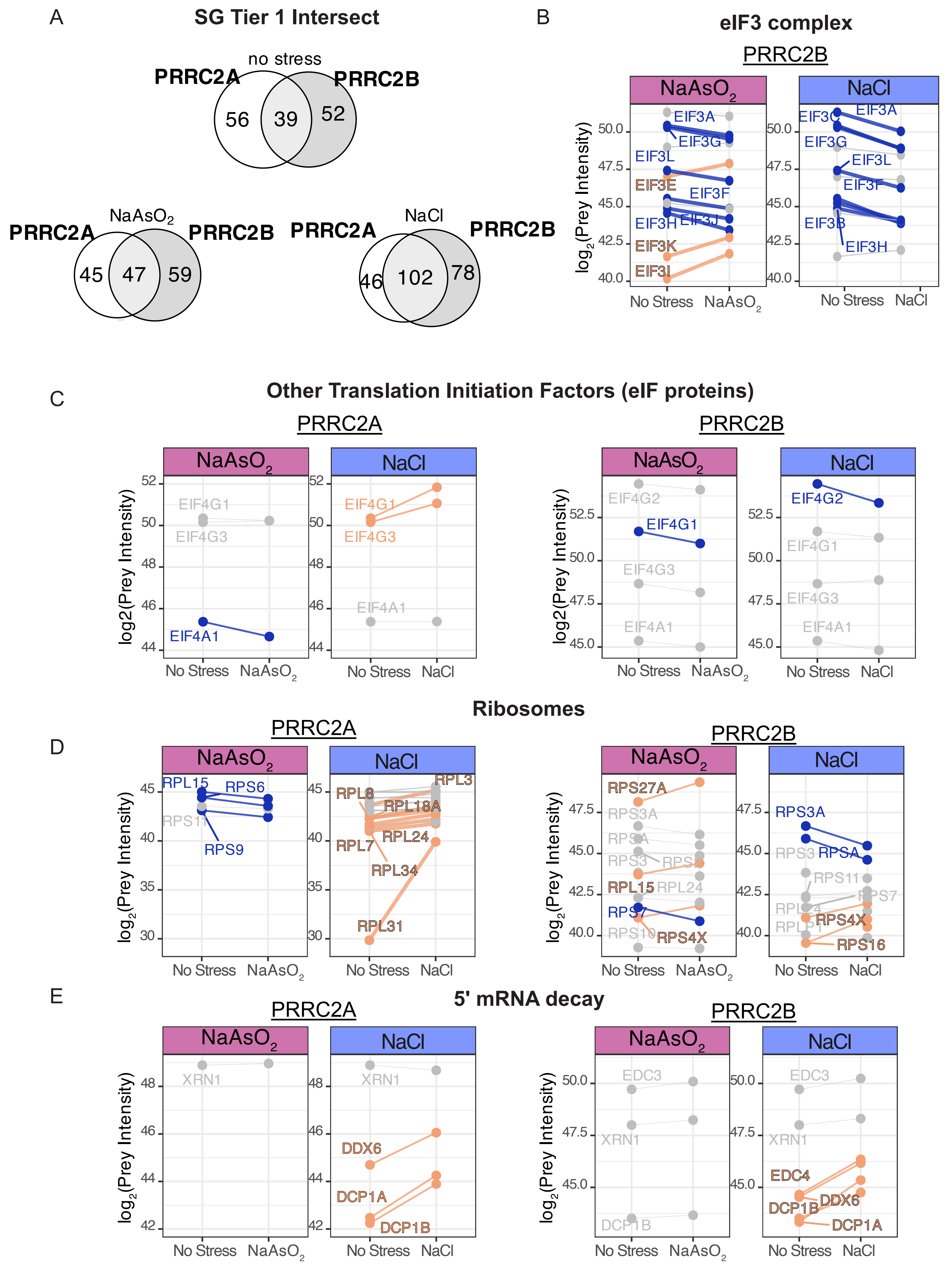

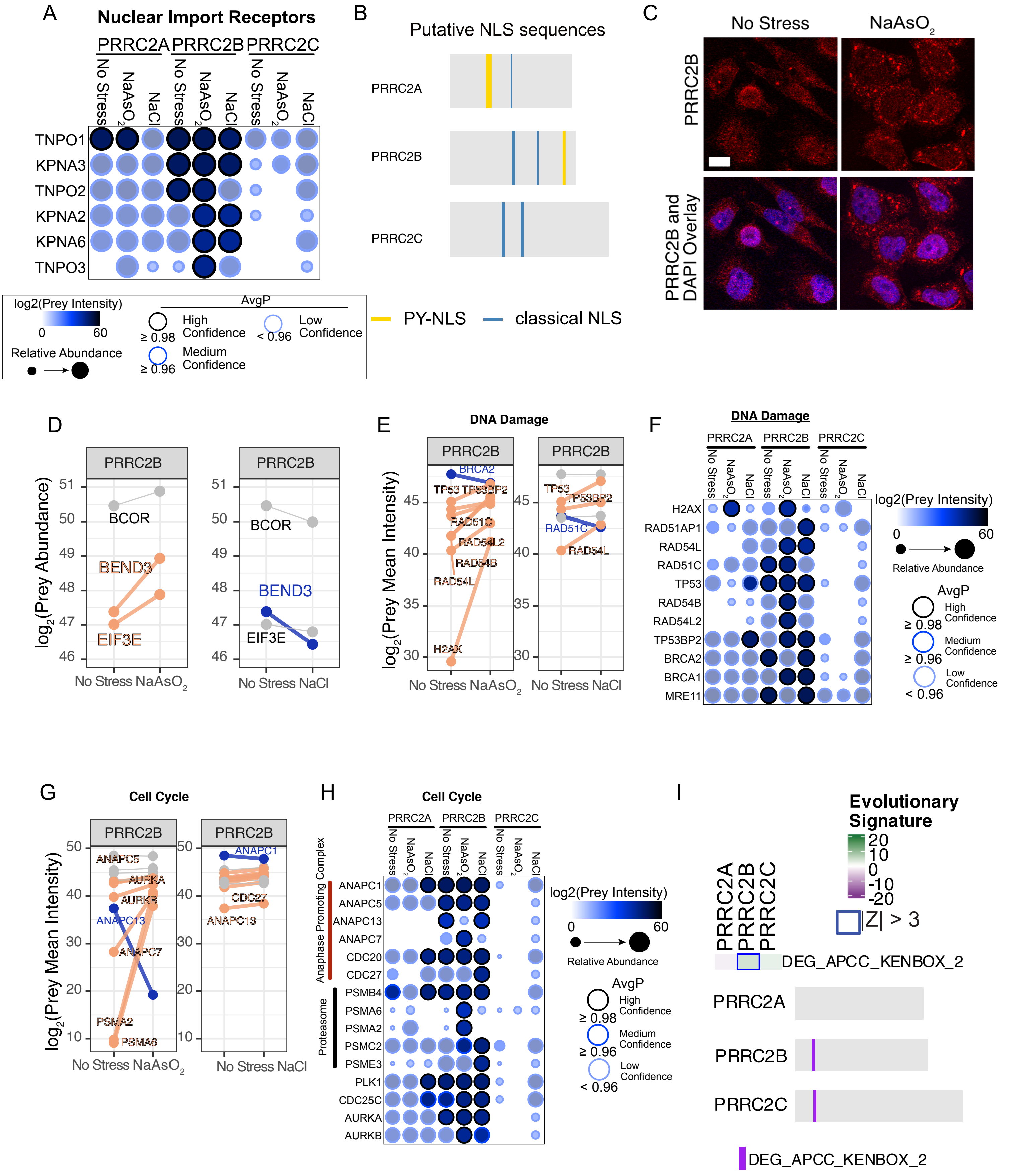

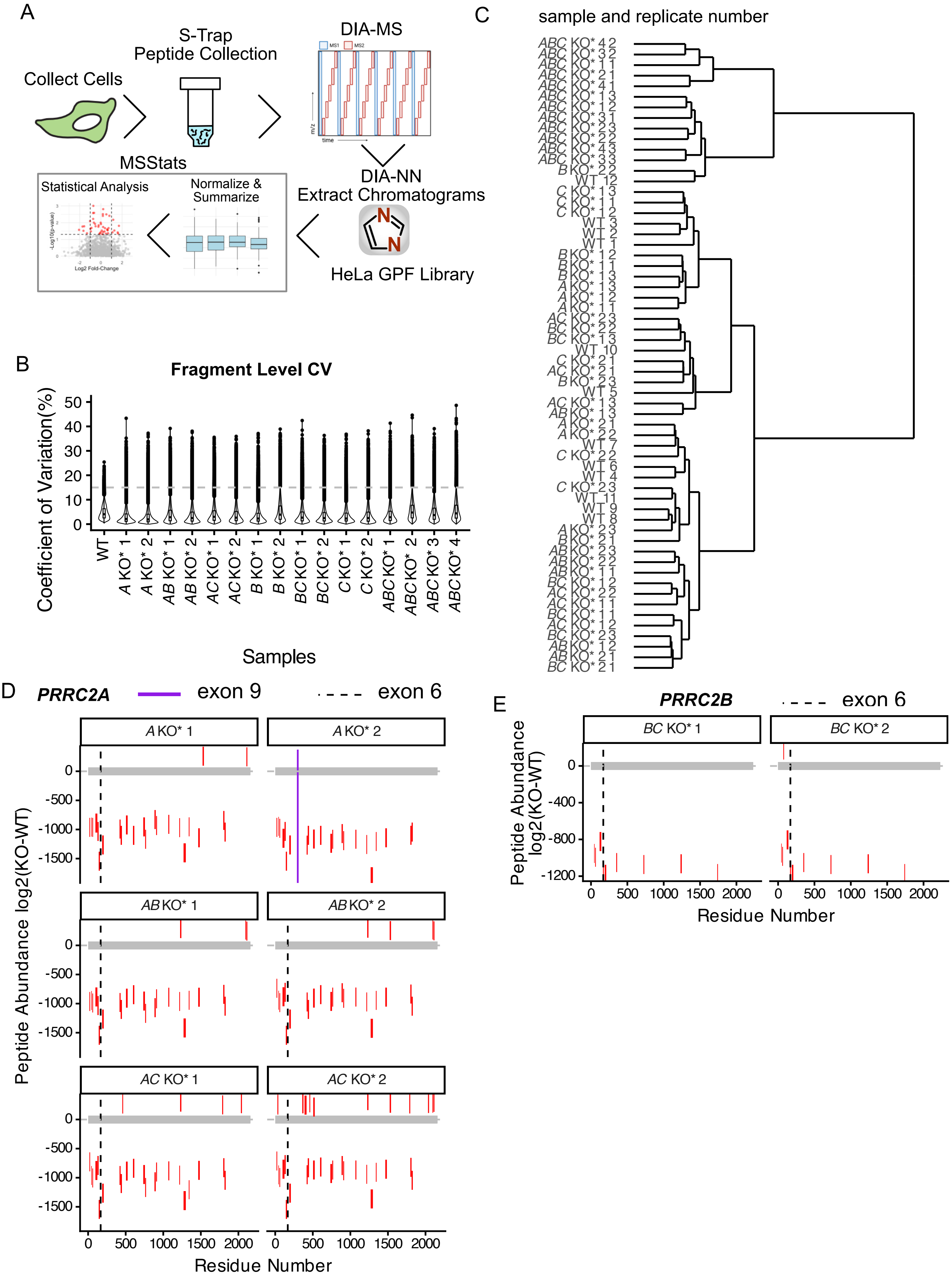

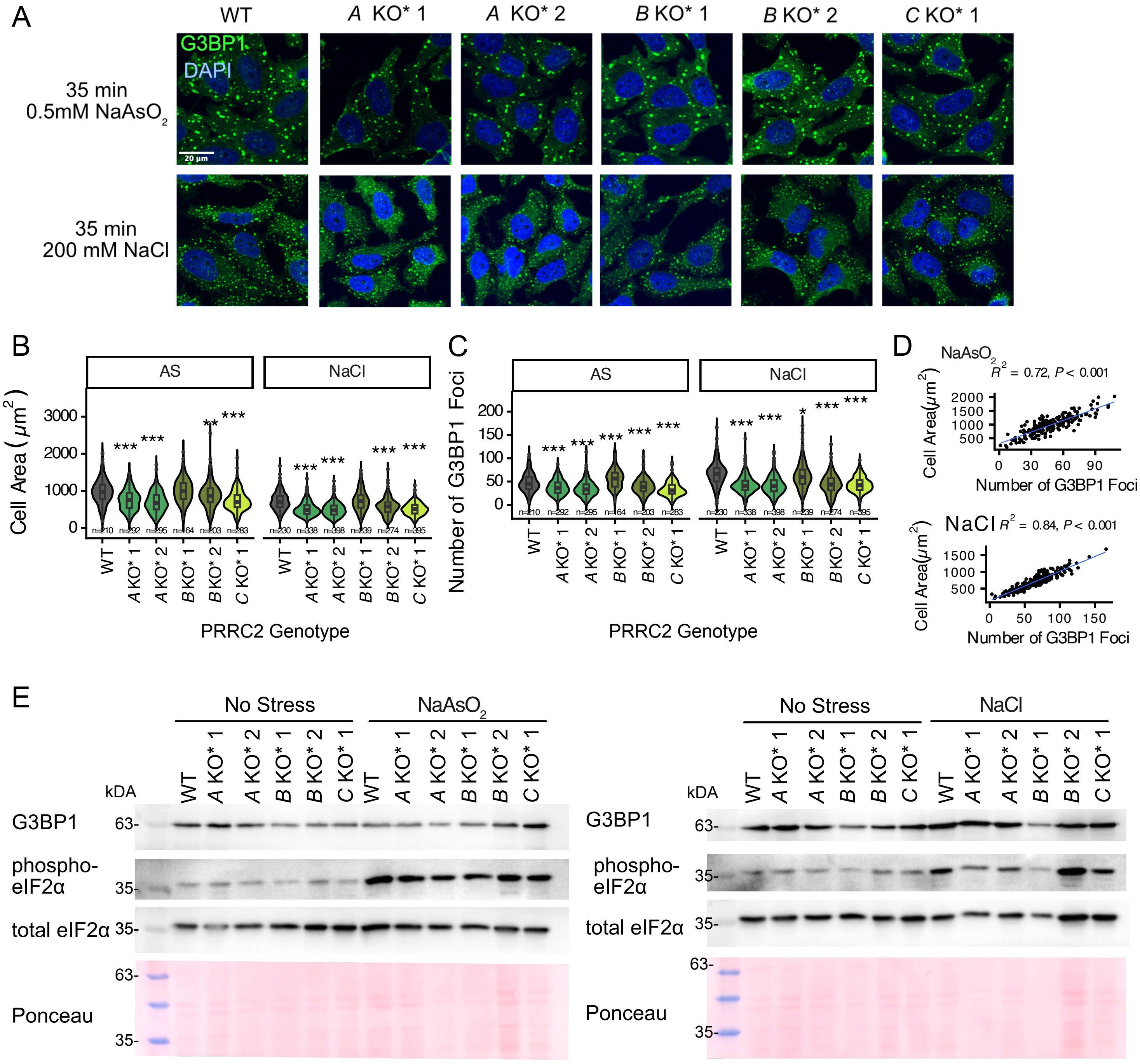

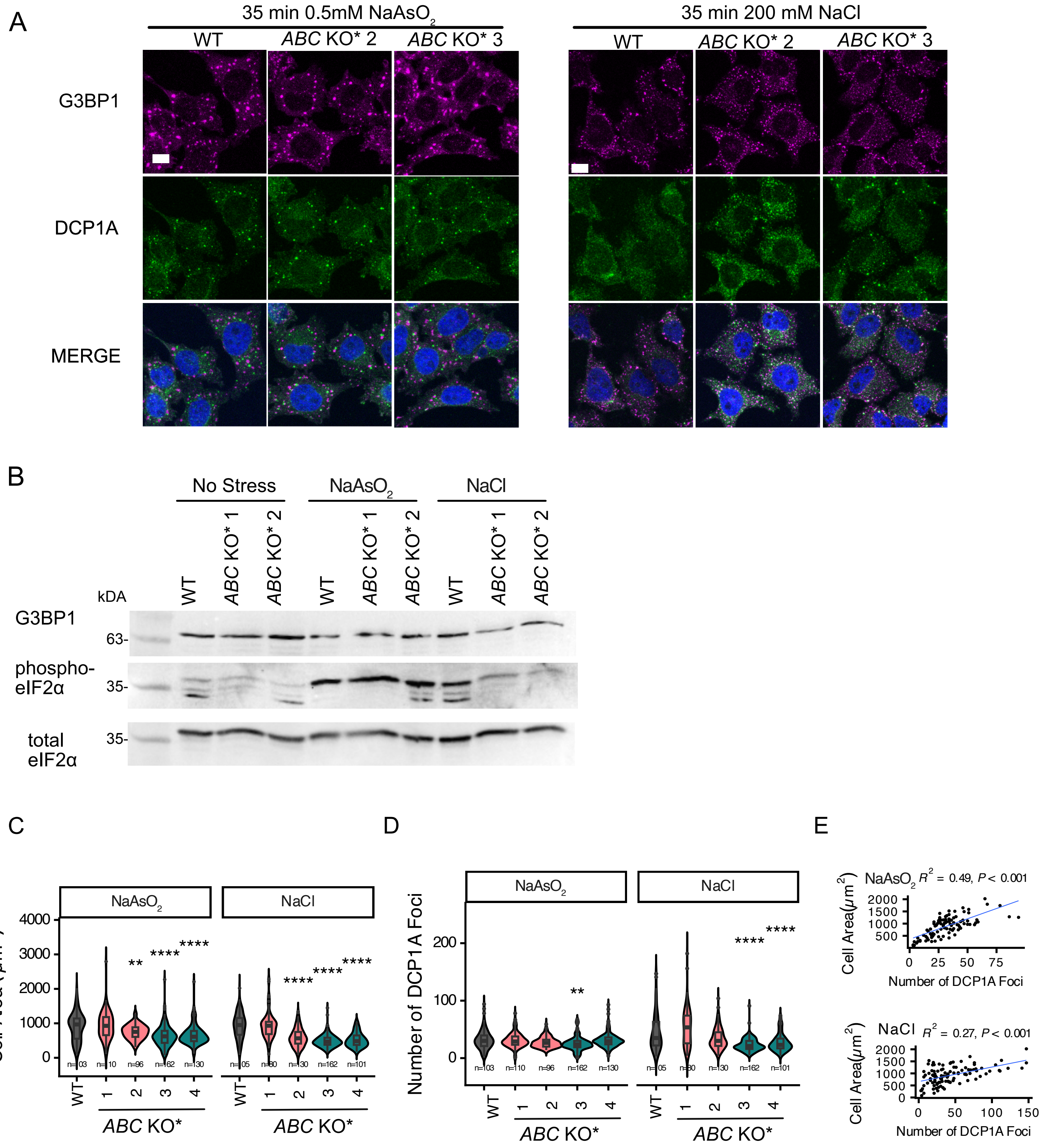

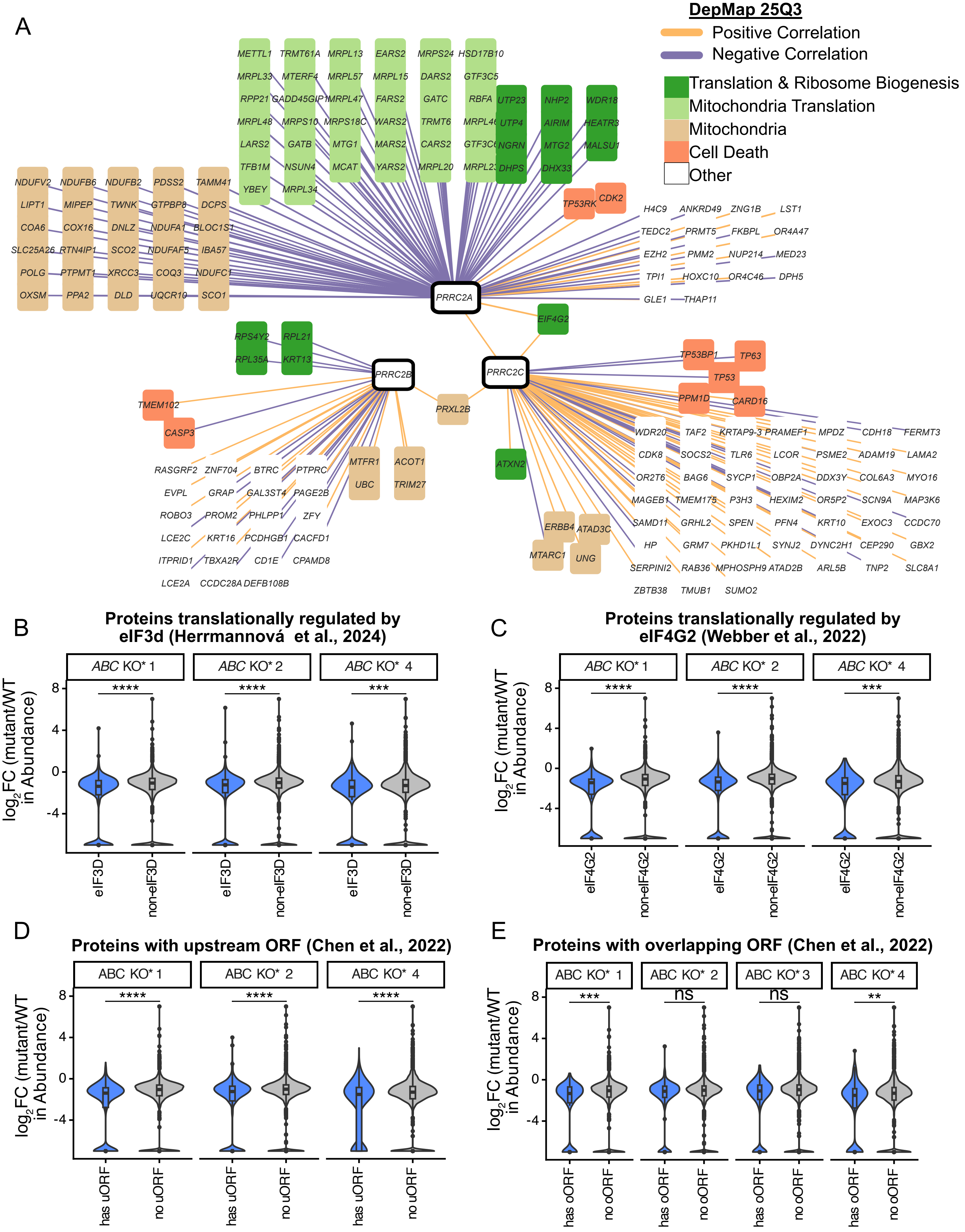

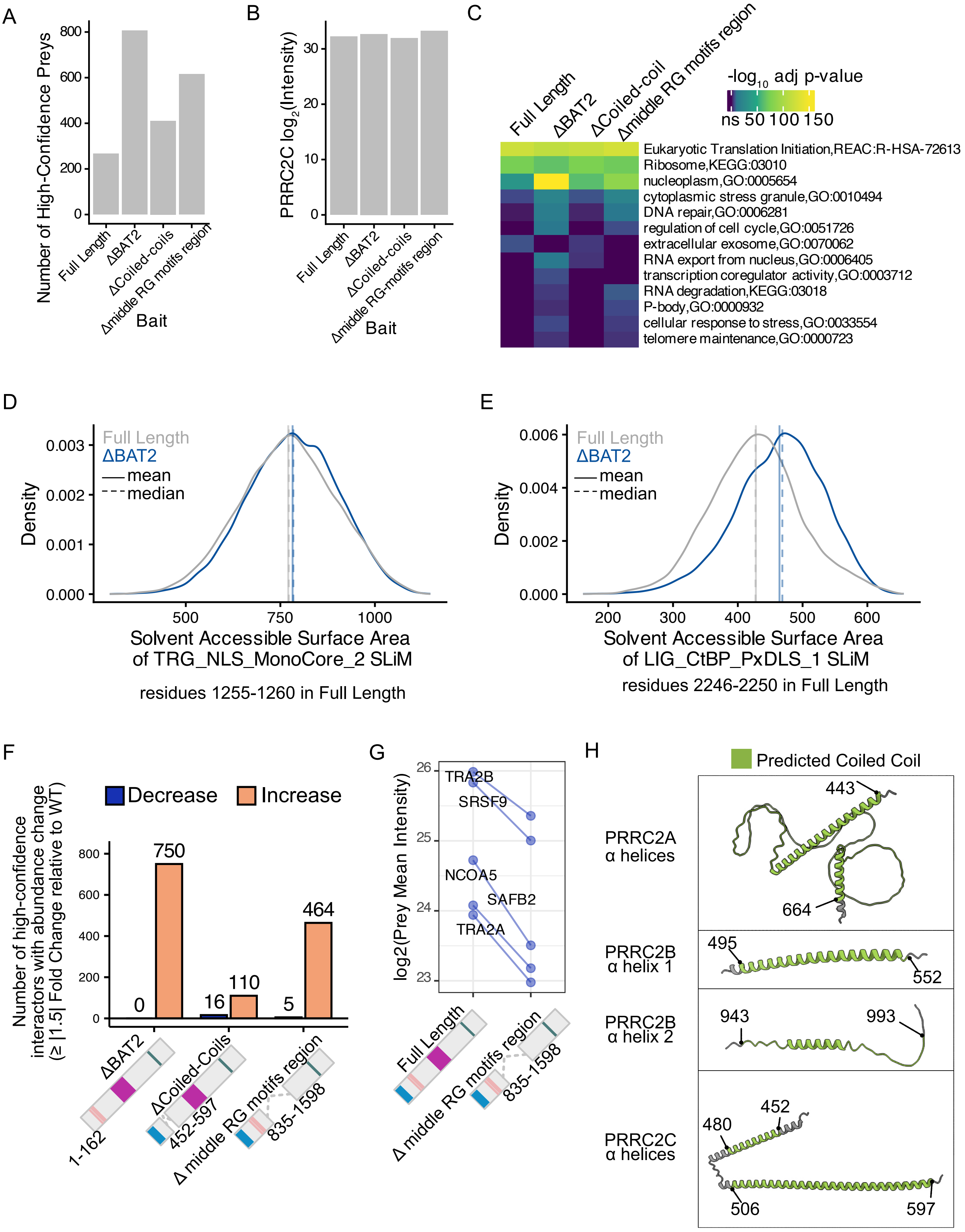
